# A novel description of the network dynamics underpinning working memory

**DOI:** 10.1101/2023.01.20.524895

**Authors:** Chiara Rossi, Diego Vidaurre, Lars Costers, Fahimeh Akbarian, Mark Woolrich, Guy Nagels, Jeroen Van Schependom

## Abstract

Working memory (WM) plays a central role in cognition, prompting neuroscientists to investigate its functional and structural substrates. The WM dynamic recruits large-scale frequency-specific brain networks that unfold over a few milliseconds – this complexity challenges traditional neuroimaging analyses. In this study, we unravel the WM network dynamics in an unsupervised, data-driven way, applying the time delay embedded-hidden Markov model (TDE-HMM). We acquired MEG data from 38 healthy subjects performing an n-back working memory task. The TDE-HMM model inferred four task-specific states with each unique temporal (activation), spectral (phase-coherence connections), and spatial (power spectral density distribution) profiles. A theta frontoparietal state performs executive functions, an alpha temporo-occipital state maintains the information, and a broad-band and spatially complex state with an M300 temporal profile leads the retrieval process and motor response. The HMM states can be straightforwardly interpreted within the neuropsychological multi-component model of WM, significantly improving the comprehensive description of WM.

**Highlights:**

- Working memory recruits different frequency-specific brain networks that wax and wane at a millisecond scale.
- Through the time-delay embedded hidden (TDE-HMM) we are able to extract data-driven functional networks with unique spatial, spectral, and temporal profiles.
- We demonstrate the existence of four task-specific brain networks that can be interpreted within the well-known Baddeley’s multicomponent model of working memory.
- This novel WM description unveils new features that will lead to a more in-depth characterization of cognitive processes in MEG data.

## 1 Introduction

Working memory (WM) is a higher-order cognitive function that entails the temporary maintenance and manipulation of a limited amount of information^1^. Any cognitive task, from language comprehension to mathematical reasoning, relies on WM, making it one of the most studied cognitive domains^2,3^. Whilst neuropsychology has described WM with conceptual models, neuroscience has investigated the functional and structural substrates of the brain dynamics underpinning WM^4^.

In the most widely accepted neuropsychological model, Baddeley describes WM as a multi-component system in which each unit comprises different WM subprocesses: the central executive performs encoding, whereas the phonological loop (for verbal stimuli) and visuospatial sketchpad (for spatial stimuli) store the memory items^1,4,5^. In their reviews, Baddeley and Chia et al. have mapped this multi-component model onto the brain. They linked the WM units to specific brain regions yielded from neuroimaging studies, in particular, based on fMRI (functional magnetic research imaging) research^1,4^.

fMRI studies have revealed the large-scale brain networks that characteristically activate during WM tasks, namely the frontoparietal network and the default mode network (DMN)^6–8^. However, the resulting maps of brain functioning supporting WM are static and simplistic. One reason rests on the coarse temporal resolution of fMRI data, which cannot access cognitive processes unfolding over a few milliseconds^9^. Secondly, fMRI studies do not provide any information on the spectral dimension of neural activity.

Electrophysiological studies considering magneto-and electro-encephalographic (M/EEG) data overcome these shortcomings. Traditionally, event-related potentials (ERPs) extracted from EEG data have described the millisecond temporal evolution of region-specific WM activity, identifying characteristic traits such as the P300^10–13^. The morphology (amplitude and latency) of this peak is altered in cognitively impaired subjects, making the P300 a cognitive marker and descriptor for WM^11,13,14^. Instead, time-frequency analyses of M/EEG data have explored the frequency-specific power activation of the neural populations over time in a single region of interest^15–17^. Among the functionally relevant rhythms in WM, the excitatory theta (4-8 Hz) and the inhibitory alpha (8-12 Hz) activities are noteworthy. The former arises in the prefrontal regions during executive control functions, whereas the latter is associated with maintenance in temporal and occipital regions^12^.

So far, we have observed that the brain functioning underpinning WM has been explored in the spatial (fMRI studies), temporal, and spectral (ERPs and time-frequency studies) dimensions. However, findings of different studies are difficult to merge, and studies that focus only on one or two of these dimensions or severely sacrifice temporal resolution offer an incomplete view of WM. Therefore, the WM literature lacks a study investigating, in a comprehensive way, the WM frequency-specific network dynamics that unfold over a few milliseconds. Merging this with the neuropsychological understanding of WM would remarkably improve the comprehension of the brain functioning supporting WM.

A novel technique that simultaneously accesses the three dimensions (space, time, and frequency) of brain dynamics is the time-delay embedded hidden Markov model (TDE-HMM). This Bayesian model depicts the data as emerging from the activation of a set of latent states, representing power and coherence across brain regions. The HMM technique was already applied to analyse MEG resting-state data in healthy^18–20^ and pathological^21^ conditions, as well as task data^22,23^. The HMM data-driven brain states resemble the fMRI resting-state networks (RSNs) in the spatial configuration. However, they present additional temporal (time of activation) and spectral (phase-coherence across brain regions) profiles -these last spanning a set of data-driven frequency bands -, providing three-dimensionality to the static fMRI functional networks.

In this study, we apply the TDE-HMM to analyse MEG data acquired during a working memory paradigm, the n-back task. The MEG data provide a millisecond temporal resolution, and the model infers four data-driven task-relevant states describing the WM dynamics without compromising the temporal resolution of the data. The HMM states are easily linked to the different parts of the multi-component model. A theta frontoparietal state performs executive functions, a temporal-occipital state with suppression of alpha inhibitory rhythm stores memory items, and a broad-band and spatially complex state with an M300 temporal profile leads the retrieval and decision-making processes to motor response. These findings are consistent with the described WM literature, unveiling new traits of the WM dynamics, such as the M300 state. Altogether, our study provides, for the first time, a 360° description of the time, space, and frequency domains of the fast transient networks underpinning working memory.

## 2 Methods

### 2.1 Participants

We included 38 healthy subjects with normal to corrected vision. In Table 1, we present the demographics of the dataset. All participants signed informed consent, and the study was approved by the ethics committees of the National MS Center Melsbroek and the University Hospital Brussels (Commissie Medische Ethiek UZ Brussel, B.U.N. 143201423263, 2015/11).

**Table 1.** Demographic data of the participants. We report the female and male groups, separately; for each group, the age and education are expressed as mean and standard deviation (std). The p values result from a one-way ANOVA test for age and education separately, considering gender as the grouping variable, with levels ‘Male’ and ‘Female’.

|  | Number of Subjects | Age - years (mean± std) | Education - years (mean± std) |
| --- | --- | --- | --- |
| Male | 15 | 49.4± 6.9 | 14.9± 3.1 |
| Female | 23 | 42.7± 11 | 14.7± 3.3 |
| p |  | 0.05 | 0.8 |

To test the demographics of the dataset (age and education), we ran a one-way ANOVA test, considering gender as the grouping variable.

### 2.2 MEG and MRI Data Acquisition

Every subject underwent an MEG and MRI acquisition. The MEG data were acquired at the CUB Hôpital Erasme (Brussels, Belgium) with two scanners: the Neuromag VectorViewTM system (13 subjects), and then the updated NeuromagTM TRIUX system (MEGIN Oy, Croton Healthcare, Helsinki, Finland) (25 subjects). Both MEG systems consist of 102 triplets of sensors, each including 2 planar-gradiometers and 1 magnetometer. The device lays in a lightweight magnetically shielded room (MSR, MaxshieldTM, MEGIN Oy, Croton Healthcare, Helsinki, Finland). Prior to the MEG recording, the shape of the subject’s scalp was recorded using an electromagnetic tracker (Fastrak, Polhemus, Colchester, Vermont): more than 400 points were traced over the whole scalp, nose, and face, in addition to the 3 fiducial points (nasion, left and right preauricular). During the MEG acquisition, participants sat in the MEG scan with 3 coils on the mastoid, left, and right forehead to track the head’s movements, and sensors to record an electrocardiogram (ECG), and electrooculogram (EOG). The MEG signal was acquired with a sampling frequency of 1000 Hz, and a [0.1 330] Hz band-pass filter.

The MRI data were collected at the Universitair Ziekenhuis Brussel (Jette, Belgium), using a 3T Achieva scanner (Philips, Best, Netherlands). The 3D MR images were T1-weighted, and the recording parameters were TR = 4.939 ms, FA 8, 230 × 230 mm^2^ FOV, 310 sagittal slices, resulting in a 0.53 by 0.53 by 0.5 mm^3^ resolution. This image was affinely coregistered to the MNI152 atlas. The structural and functional acquisitions were collected with 5 days (median value) in between (IQR 2-10 days).

### 2.3 Task design

All participants performed an n-back task during the MEG recording. The n-back task consists in showing a sequence of letters, and the subject is instructed to respond to a target letter by pressing a button with the right hand. In the 0-back condition, the letter X is the target; during the 1-back condition, the target is any letter that coincides with the preceding one; at last, in the 2-back condition, the target is any letter that coincides with the one shown 2 letters before. The rest of the shown letters are considered distractors. Figure 1 visually explains the task, also providing the timing information.

**Figure 1.**
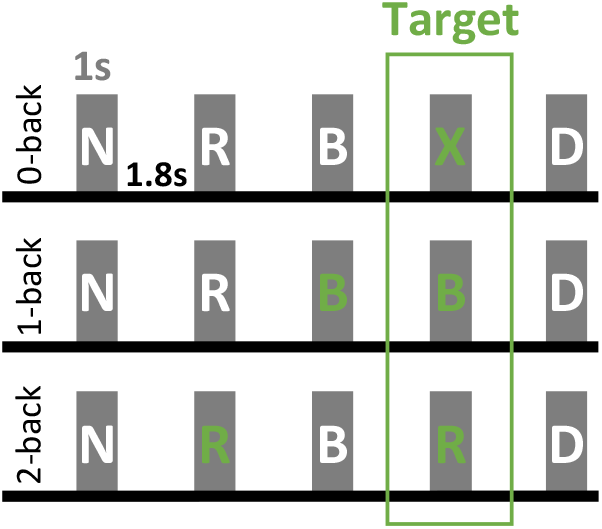
Graphic representation of a visual-verbal n-back task. Every letter is displayed for 1 second, and the inter-trial period is 1.8 sec. Every green letter represents the template to which every target is matched, and the target letters – for which subjects must press a button -are in the green rectangle.

The experimental setup consisted of a screen located 72 cm from the MEG helmet. Letters projected on the screen fit within a 6×6,5 cm area. A photodiode below the screen detected the stimulus onset. The reaction time was then computed as the time between the stimulus onset detected by the photodiode and the moment when the subject pressed the button.

Participants went on a training session before the actual recording to verify whether they understood the task. Twelve blocks of 20 letters (stimuli) each were presented pseudo-randomly, four for each paradigm condition. The total number of target trials is 25, 23, and 28, for the 0, 1, and 2-back conditiona, respectively.

### 2.4 Data pre-processing, parcellation, and sign flipping

#### Data preprocessing

The entire analysis was developed in MATLAB 2020b, and the data preprocessing was carried out following the MEG analysis pipeline proposed in ref^22^. The pipeline uses the Oxford’s Software Library and builds upon SPM12 (Welcome Trust Centre for Neuroimaging, University College London) and Fieldtrip^24^. First, we coregistered the MEG data to the T1 MR image of the same subject, applying the RHINO algorithm. Here, we used the subject-specific fiducial points acquired with the Polhemus tracker, as explained in the previous paragraph, to minimize the coregistration errors. Next, we downsampled the MEG data to 250 Hz and applied a band-pass filter [1, 45] Hz -Butterworth IIR filter of order 5 with zero-phase forward and reverse filter, the instability is solved by reducing the order of the filter - to discard the high and low-frequency noise. We also included a notch filter around 50 Hz to further remove the power line effect. We performed artefact rejections in two steps. First, we identified bad data segments for each subject separately. Segments of one second with an outlier standard deviation were labeled as “bad” and subsequently discarded. Next, we applied the AFRICA algorithm that decomposes the data into 62 independent components (ICA) and removes those that correlate with ECG and/or EOG (r>0,5). In the last step, we visually examined the data to verify that all major artefacts were removed.

Because the MEG data are acquired by magnetometers and planar gradiometers, the data variance must be normalized across sensor types. The data relating to each sensor are decomposed in eigenvalues and normalized by the smallest eigenvalue^25^. Afterward, we applied the linearly constrained minimum variance (LCMV) beamforming algorithm to reconstruct the source space^25^. The source reconstruction was based on a single-shell forward model in MNI space with a projection on a 5 mm dipole grid.

#### Parcellation

To parcel the source-reconstructed data, we used a 42 parcels atlas, used before in^21,23^. This data-driven parcellation was extracted using ICA decomposition of fMRI data from the Human Connectome Project. The atlas includes only cortical regions. The time course for each ROI was extracted as the first principal component across the voxels’ time series. As beamforming may lead to signal leakage between regions, we orthogonalized the parcels’ time series by multivariate symmetric leakage correction ^26^. This conservative approach discards signal components that are very close to zero lag; nonetheless, this step assures that the following connectivity analysis is not affected by neighboring signal leakages.

#### Sign flipping

After beamforming, the sign of the resulting time course is not uniquely defined. The sign of the dipoles is arbitrarily assigned, and this hinders the analysis across subjects and across brain regions. Therefore, we applied the sign-flipping algorithm presented by Vidaurre et al.^23^.

### 2.5 Time delay embedded-hidden Markov model (TDE-HMM)

The hidden Markov model rests on the idea that the observed data (MEG) emerge from a set of hidden underlying states (brain networks). This model assumes that the probability of activation of a state at time t depends only on the empirical data at time t and the latent state at time t-1 (order t-1).

Specifically, we implemented the time-delay embedded HMM (TDE-HMM), as described by^18^. This model takes into consideration the conduction delay characterizing the communication across brain regions, by adding a lagged version of the data to the original matrix (ROIs*time points). This is referred to as embedding. We lagged the data considering a [-7 : +7] interval of time points (14 LagPoints), that corresponds to a time window of 60 ms. This embedded matrix was then reduced applying PCA and considering 2*ROIs = 84 principal components (as recommended in^20^). A Gaussian distribution was then defined on the embedded space. The parameters of the state Gaussian distributions and the state dynamics are computed by stochastic Bayesian inference, as described in ref^23^.

Eventually, the two model outputs that we considered are the posterior probabilities and the Viterbi path. The posterior probabilities correspond to the states’ time courses, providing the probability that each state activates at each time point. In the Viterbi path, each time point is categorically (instead of probabilistically) associated with only one state.

The discrete number of states must be defined as apriori. We ran different inferences with 4, 6, and 8 states to explore how the model would describe the dynamics of the data in the different scenarios.

### 2.6 Epoching and GLM statistics

We performed an event-related field analysis starting from the state time courses, following the methods previously explained by^22^. The time course of each state is epoched with respect to the stimulus onset, taking an epoch of 1400 ms long, [-200 1200] ms. Each trial was baseline corrected considering the pre-stimulus window [-200 -30] ms. Afterward, we ran a two-level generalized linear model (GLM) to investigate the states’ activation pattern. The GLM design matrix consisted of 7 regressors, the constant regressors (mean activity over all task conditions), and the 6 paradigm conditions: 0 back target, 1 back target, 2 back target, 0 back distractor, 1 back distractor, 2 back distractor. Additional contrast regressors were used to evaluate the effect of response (target vs distractor) and working memory load (0, 1, and 2 back) on the states’ activation patterns. The GLM first computed the contrast of parameter estimates (COPEs) for each subject (first level), and afterward, the mean COPEs per subject were fitted across subjects (second level) to evaluate the probability that a given state is active at a certain point in time. At the group level, we tested whether the mean COPE at each timepoint was significantly different from zero with a non-parametric permutation test (1e4 permutations). The significance threshold was set to 97.5% of the null distribution, and the multiple comparison problem was overcome by taking the maximum statistic across time and states. For additional details on this methodology, we refer to^22^.

### 2.7 Power spectral densities (PSD) and Phase-coupling Connections

The power spectral density is estimated by applying a non-parametric multitaper to the original data weighted by the posterior probability of each state in the broad frequency band 1-40 Hz; for more details on the method refer to^18,20^. This multitaper is applied state by state and the PSD is estimated for each subject and state separately. The same approach is also applied to extract the phase-coherence between pairs of regions over all the parcellation for each state, forming the state-specific phase-coupling network.

### 2.8 Spectral modes

The spectral information of each state is extracted in 4 spectral modes that correspond to 4 frequency bands in the broad band (1-40 Hz). These spectral modes are defined in a fully data-driven way. Starting from the phase-coupling matrix of each state and subject, a non-negative matrix factorization (NNMF) is run across all nodes, connections, subjects, and states, as explained by Quinn et al.^22^. We only consider the first three spectral modes, as the fourth one is extracted to filter the high-frequency noisy components, and then we discard it^27^. The NNMF provided the profiles of the four spectral modes, the state-and subject-specific phase-coherence connections and PSD distributions for each spectral mode. The PSD values are normalized (z-score) for visualization. The strongest connections in the phase-coupling networks are identified by Gaussian Mixture model (GMM).

## 3 Results

### 3.1 Participants

Regarding the demographics of the dataset, the male and female groups do not significantly differ in age (one-way ANOVA test, F = 4.3, p_val = 0.05) or education (one-way ANOVA test, F = 0.05, p_val = 0.8) (ref. Table 1). We report the mean reaction times (mean RTs) and the accuracy of response per paradigm condition in Figures S1 and S2 of the supplementary materials.

### 3.2 Number of States, model reliability, and replicability

We ran the model by setting the number of states to infer at 4, 6, and 8. Increasing the number of states allows us to explain more data variance; however, we also increase the probability of redundant information shared across states^28^. We used the task-evoked activation pattern from the GLM analysis (the state average response activation as shown in Figure 2, plot A) to visually assess the inferences. Considering the WM literature, we expected at least three relevant states, i.e. a frontal/frontoparietal state, a state associated with a motor response, and a state capturing the alpha dynamic. We observed that the four states inference could not always capture the expected brain dynamic traits. Comparing the 6 and 8 states inferences, we chose 6 as the optimal number of states as we noticed that, in the 8 states inference, some states were overlapping in PSD maps and task-evoked activation patterns.

**Figure 2.**
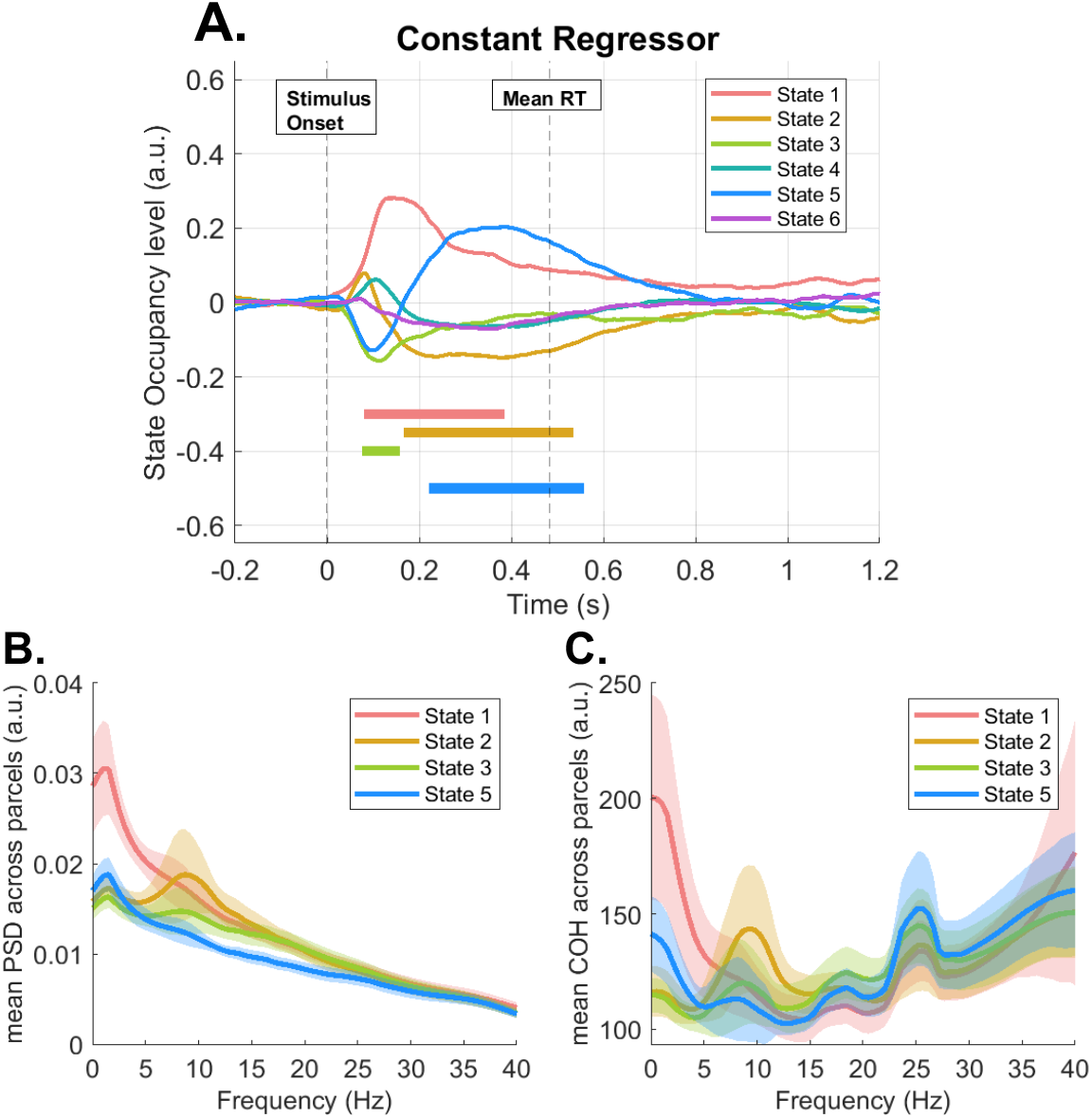
States description: (A) state average response across conditions, (B) distribution of the state-wise power spectral density (PSD) (C) distribution of the state-wise mean coherence (COH) – PSD and COH are averaged across subjects and regions/pair of regions. The straight lines at the bottom indicate the time points where the state of the same color is significantly activated or deactivated; permutation test, significance level 0.025. The significantly activated, task-relevant, states are state 1, state 2, state 3, and state 5. State 1 focuses mostly on low frequencies and state 2 presents a peak around 10 Hz, both for power (A) and coherence (B) distributions. State 3, instead, shows only slightly higher power and coherence between 15 and 25 Hz. Eventually, state 5 is characterized by high coherence around 25 Hz, but flat power distribution over the frequency range.

We assessed the model’s reliability and replicability by running the model 4 times (always with 6 states). We visually compare the inferred states, as suggested by^22^, and concluded that the model could consistently infer a similar set of states. As an additional reliability test, we computed the temporal characteristics (life time, interval time, and fractional occupancy) of the states over the whole inference. If the results significantly differed from what was reported in the HMM literature, this would signify a bad inference or a problem in the model inference. We report this analysis in the supplementary materials (section 2, Figure S3). The states’ average life time is 73 ms, the average fractional occupancy is 18%, and the average interval time is 500 ms. These results are consistent with the HMM literature on resting-state and task data^22,28^.

### 3.3 ERF analysis of states’ time courses

Figure 2 (plot A) shows the average activation across all paradigm conditions of the 6 different HMM states during the n-back task. From this task-evoked activation plot, we identify the task-relevant state as those that are significantly activated or deactivated: states 1, 2, 3 and 5 (p < 0.025). The non-task-specific states are reported in the supplementary materials (Figures S7 and S9); they are associated with resting state and baseline activity^22^.

### 3.4 Spectral modes

Figure 3 reports the three spectral modes in which we extracted the state-specific spatial and connectivity networks: spectral mode 1 includes the low frequencies, especially the conventional delta (1-4 Hz) and theta (4-8 Hz) bands, spectral mode 2 focuses on the alpha (8-12 Hz) band, and spectral mode 3 refers to the beta (12-30 Hz) band.

**Figure 3.**
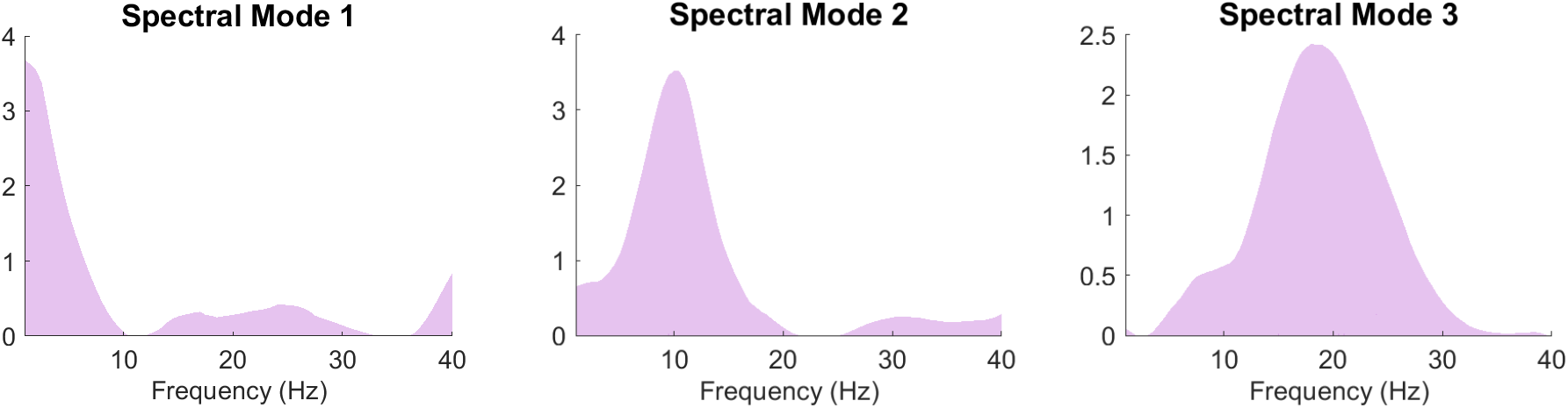
Data-driven frequency bands in which we describe the states frequency content. Spectral mode 1 is associated with the low frequencies, hence, the theta and delta conventional bands. Spectral mode 2 is associated with the alpha band, and spectral mode 3 with the beta band.

### 3.5 States Description

We propose the description of each state as composed of (1) the time course of activation (state task-evoked response) for all the paradigm conditions, (2) the mean z-score power spectral density (PSD) distribution over the brain, and (3) the phase-coupling network over the brain.

#### State 1 – The frontal executive state

State 1, depicted in Figure 4.1, represents an early low-frequency frontal network. This state is significantly activated between 150 and 350 ms (peak at 200 ms) peristimulus (PST) in all paradigm conditions. It presents a low-frequency peak of both PSD (Figure 2, plot B) and coherence (Figure 2, plot C). Therefore, most of the spectral content of this state lays in spectral mode 1. The low-frequency peak of mean PSD appears in the right and left orbitofrontal cortices (OFCs), the medial prefrontal cortex, the right and left anterior temporal cortices, and the posterior cingulate cortex (PCC). The connectivity network shows strong phase-synchronization in the low frequencies between prefrontal and anterior temporal regions and a connection between the OFC and the PCC.

**Figure 4.**
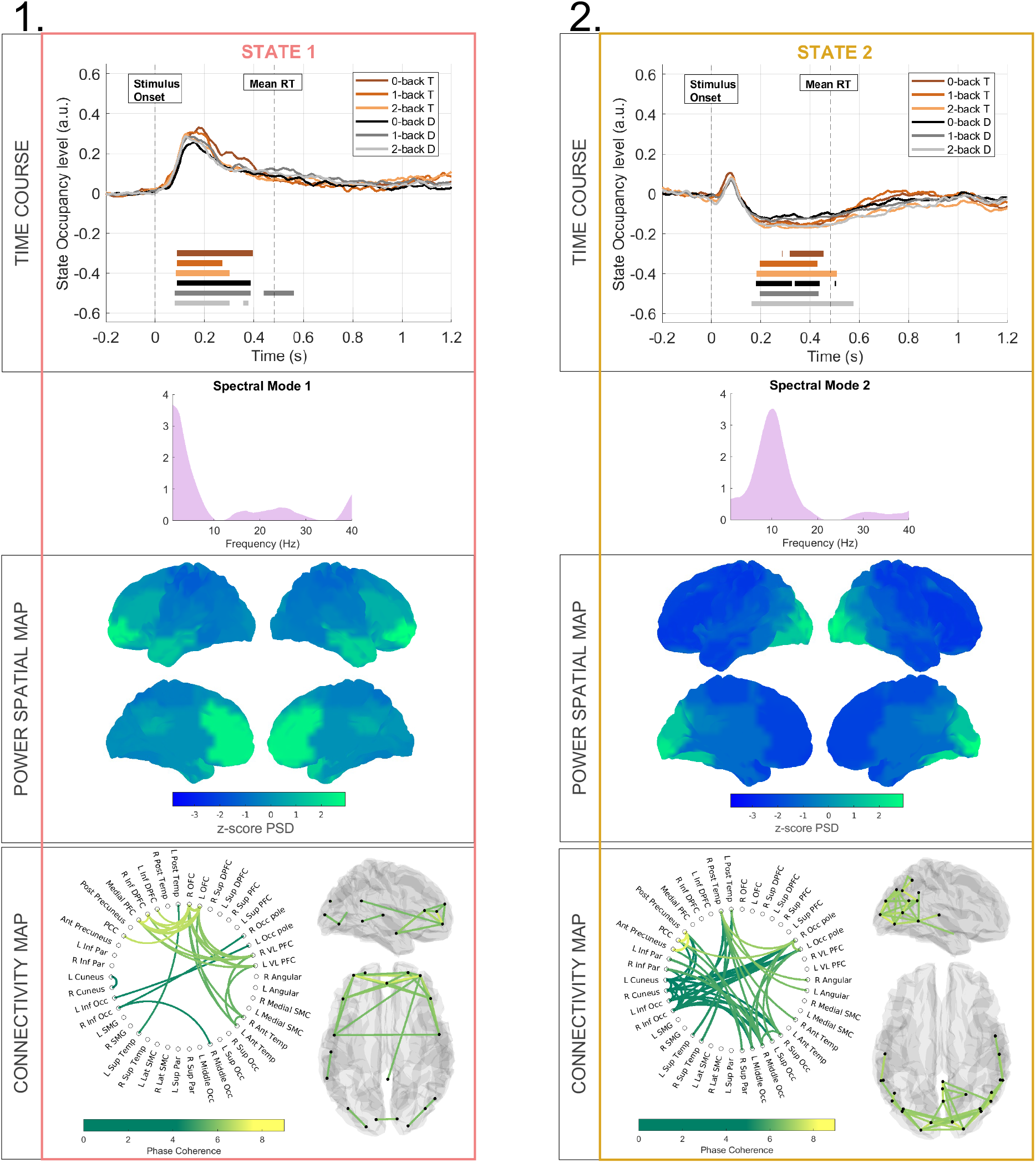
Main characteristics of state 1 (1) and state 2 (2). In each column the top plot shows the state time course (task-evoked response) resulting from the GLM analysis for the 6 paradigm conditions, separately. The second box of the column present the spectral mode associated with the state, the third box shows the z-score mean power spectral density (PSD) map, and the fourth box shows the connectivity network (brain glass on the right and circular graph on the left). Only the phase-coupling connections surviving thresholding are plotted.

#### State 2 – the occipital state

State 2, displayed in Figure 4.2, represents an occipital alpha network. After a non-significant peak of activation around 150 ms, the state task-evoked response significantly decreases between 200 and 500 ms after stimulus onset for all task conditions. The activity of this state arises primarily in spectral mode 2, as it shows a peak of mean PSD and coherence around 10 Hz (Figure 2, plots B and C). The mean PSD distribution spatially focuses on the occipital lobe, including the right and left occipital poles, the right and left inferior and superior occipital cortices, and the inferior parietal cortices, including the left and right precuneus. The phase-coupling network reveals the synchronization in the alpha band of occipital regions with some parietal ones, such as the PCC, the posterior and anterior cingulate cortices, the angular gyri, and posterior temporal regions.

#### State 3 - The sensorimotor state

State 3, shown in Figure 5.1, represents a sensorimotor network. It displays a significant peak of deactivation at 100 ms peristimulus. The frequency content of this state focuses on spectral mode 3, associated with the beta band, both for the overall mean PSD and the phase-coherence, as observable in Figure 2 (plots B and C). The PSD map reveals high beta activity in the left and right supramarginal and the left and right sensorimotor cortices. The phase-coupling network shows broad connections that branch further than the regions we identified in the PSD map. The network reports a strong connection between the medial prefrontal cortex (PFC) and the left and right OFCs. Additionally, we observe intra-and inter-connections between the left and right posterior temporal regions, the left and right angular gyri, the left and right supramarginal and the left and right sensorimotor cortices.

**Figure 5.**
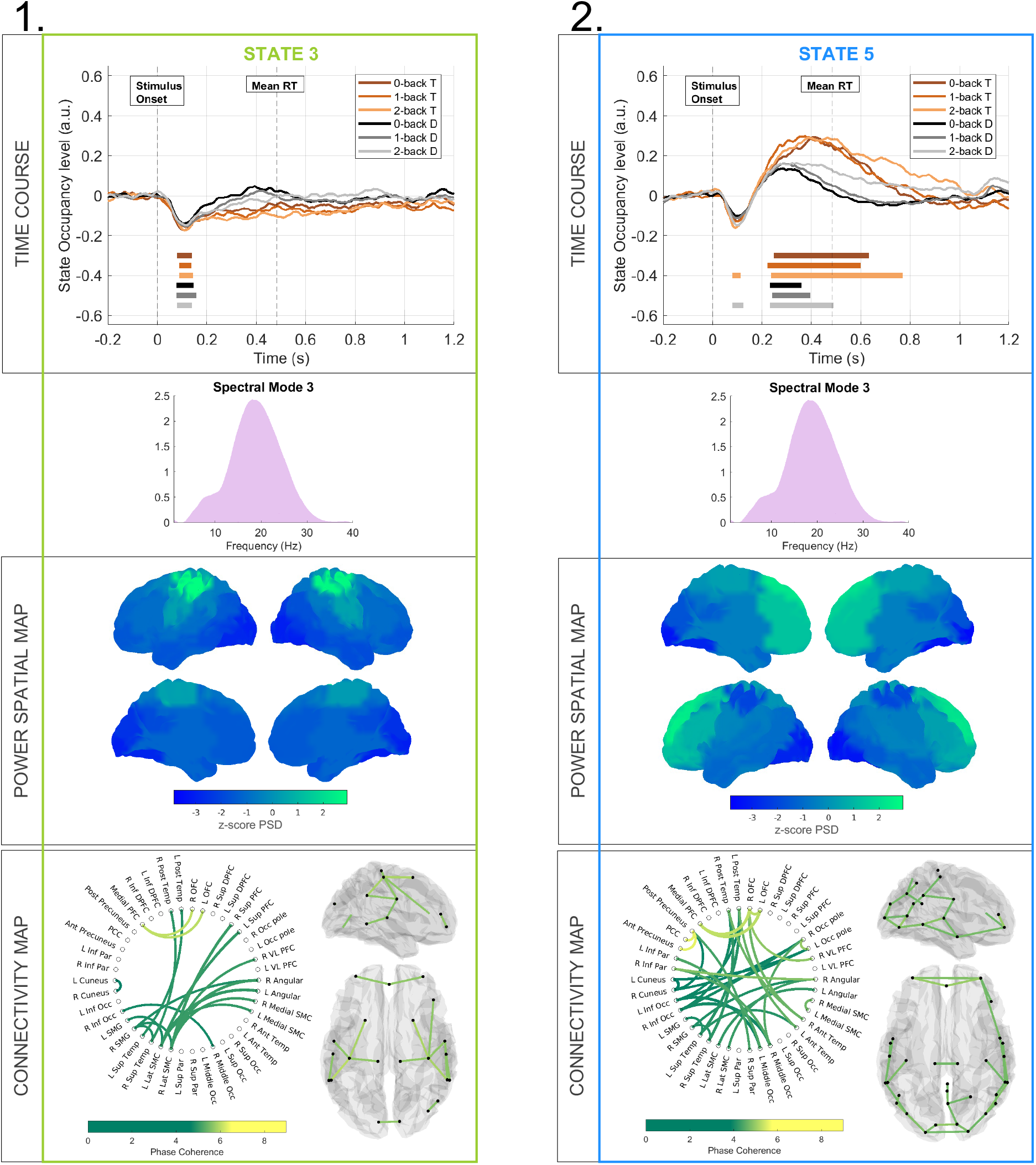
Main characteristics of state 3 (1) and state 5 (2). In each column the top plot shows the state time course (task-evoked response) resulting from the GLM analysis for the 6 paradigm conditions, separately. The second box of the column present the spectral mode associated with the state, the third box shows the z-score mean power spectral density (PSD) map, and the fourth box shows the connectivity network (brain glass on the right and circular graph on the left). Only the phase-coupling connections surviving thresholding are plotted.

#### State 5 – The M300 state

State 5, reported in Figure 5.2, displays a broad-band and spatially complex network. The task-evoked response presents a negative peak around 100 ms PST that is significant only for the 2-back conditions (target and distractor). Afterward, the activation level steadily increases, and the state is significantly activated between 225 and 500 ms PST. The state presents high mean coherence across pairs of regions and subjects around 25 Hz (Figure 2, plot C), instead, the mean PSD across subjects and regions is flattened over the frequency spectrum (Figure 2, plot B). We then associate the spectral content of state 5 with spectral mode 3. The phase-coupling associated with spectral mode 3 displays connections between the anterior and posterior precuneus and the PCC, and the right and left OFC with the medial PFC. In the same spectral mode, the mean PSD covers a wide area of the frontal cortex, including the inferior and superior dorsal PFC, the medial PFC, the left and right superior PFC, and the right and left medial sensorimotor cortex (SMC). Although we reported only the data related to spectral mode 3, this state shows a broad band activity pattern in spectral mode 2, as reported in the supplementary material (see Figure S8).

### 3.6 Response Effect – Target vs distractor

Next, in Figure 6 we observe the state task-evoked response for the contrast between target and distractor conditions. Here, we investigate which states are sensitive to changes in WM processing between distractor and target trials. The results are similar both when neglecting the task conditions (all target vs all distractor trials) and when including the task conditions (with the following contrasts of parameters estimate: 0-back TvsD, 1-back TvsD, and 2-back TvsD – we include these results in the supplementary materials, Figure S10).

**Figure 6.**
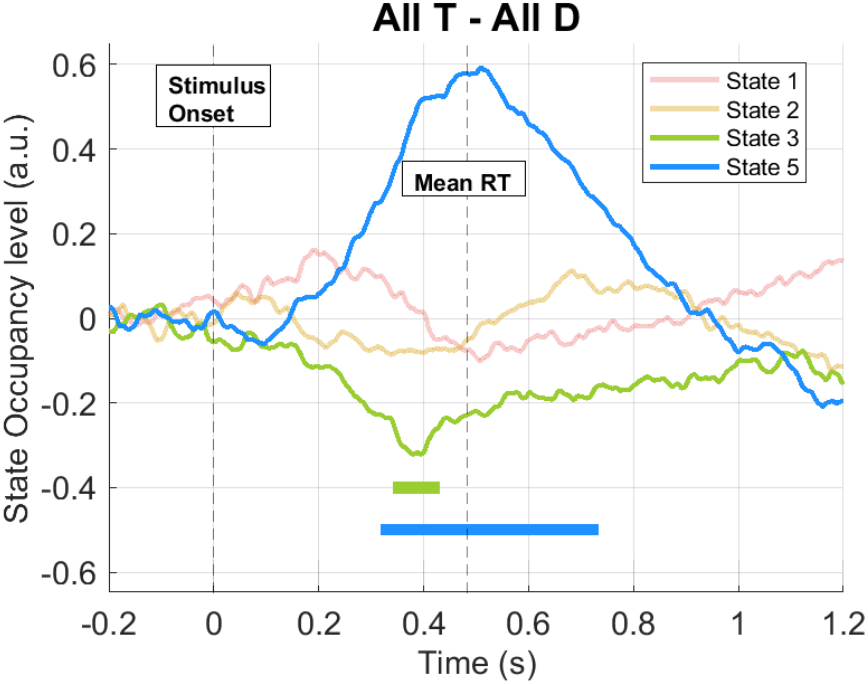
Difference of states task-evoked response between target. (together the 0, 1, and 2 back target) **and distractor** (together the 0, 1, and 2 back distractor) **trials**.

State 3, the sensorimotor state, shows a significantly decreased task-evoked response in the target compared to distractor trials around 400 ms. The change in the activation profile of the sensorimotor state is relevant since we compare targets with and without motor response. State 5 presents a significantly amplified task-evoked response in target than distractor trials between 300 ms and 700 ms.

## 4 Discussion

Working memory is a core cognitive function carried out by frequency-specific large-scale brain networks that transiently wax and wane over a few milliseconds. This study presents a novel experimental design to unravel the WM dynamic. MEG data ensure the finest temporal resolution to assess cognitive processes (ms), and the TDE-HMM extracts data-driven functional states with unique temporal, spatial and spectral profiles.

The n-back task has been extensively validated and used to investigate working memory^29,30^, as it proved to elicit a robust neural activity consistent within and between recording sessions^31^. During the short time window of an n-back epoch, the WM processes (i.e., encoding, maintenance, retrieval) are carried out almost simultaneously, with high attentional and processing demand. Therefore, the brain dynamic associated with the different WM processes results intertwined^11^.

Figure 7 reports an overview of results and discussion, showing how our work can be interpreted within the Baddeley WM framework.

**Figure 7.**
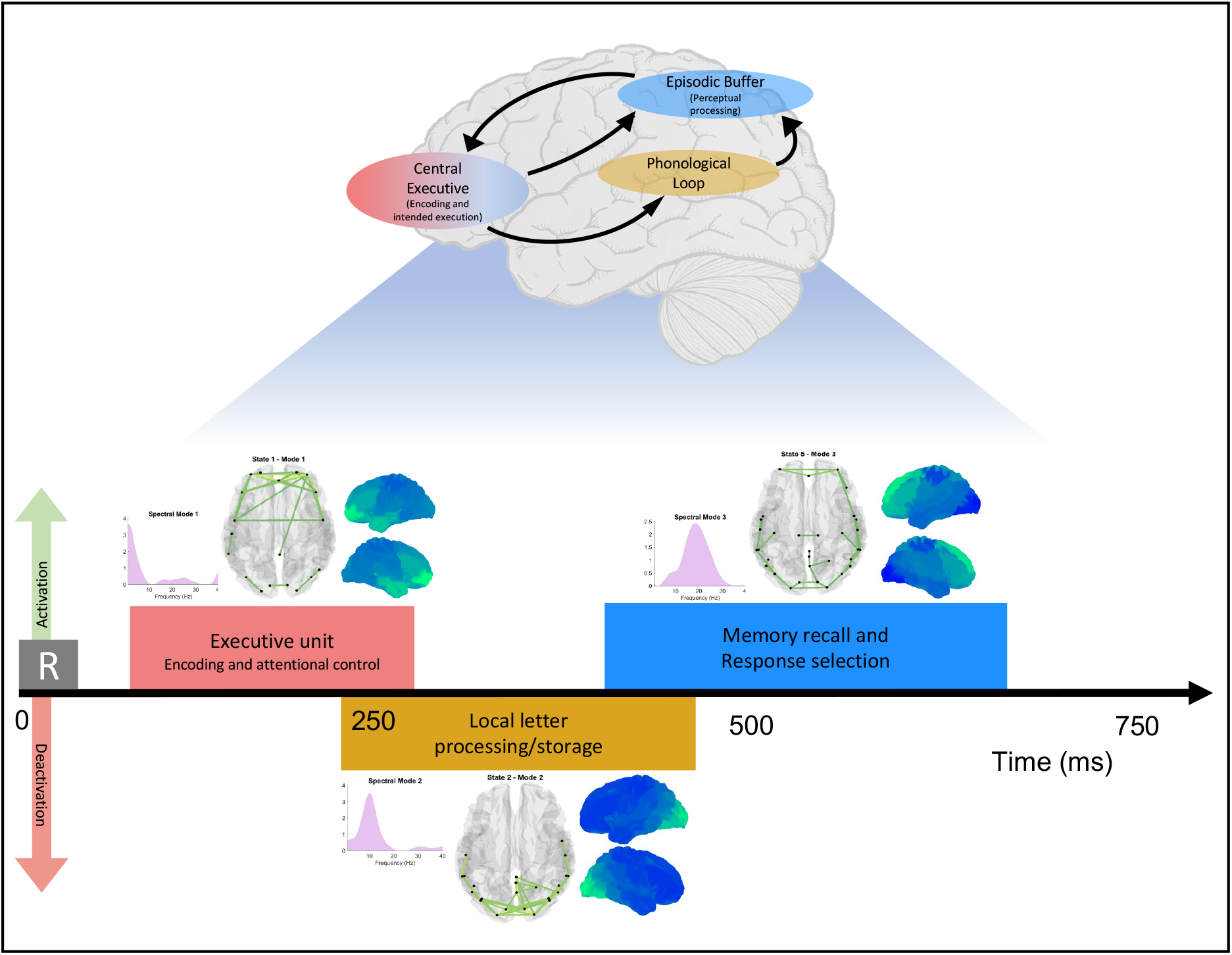
Overview of the main results of this study. On the top, we show a schematic representation of the WM multicomponent model as presented by^4,5^. On the bottom, we depict the multidimensional description of WM during an n-back task resulting from the study. The colour of each unit is associated with the colour of the state that plays the same role. We do not aim at creating a one-to-one correspondence between the data-driven results and the neuropsychological model, but rather present an integrated understanding of the WM dynamics. The HMM states provide a spatial, spectral, and temporal representation of WM dynamics, significantly improving the overall description of WM.

### State 1 – Executive control of stimulus encoding

During any WM task, the stimulus processing (‘encoding’) starts soon after the stimulus presentation. State 1 arises early after stimulus onset (see Figure 4.1) in the prefrontal cortex, suggesting that this state functions as the executive control unit. The low-frequency rhythm characterizing the spectral profile of state 1 includes the conventional theta band, which is associated with integrative and control functions during high-level cognitive processing ^15,32^. This excitatory theta activity has been repeatedly detected in prefrontal regions - consistent with the spatial map of state1 -, where the high-order representation of the stimulus occurs during WM encoding ^33–35^.

This state closely resembles the anterior DMN identified by Vidaurre et al. in resting-state data using the same TDE-HMM approach^18^. fMRI WM studies have frequently reported the DMN as actively engaged during WM encoding^6,7^.

This state is further characterized by a low-frequency frontoparietal synchronization between frontal (OFC) and parietal regions (PCC), that was previously interpreted as a volitional top-down - from high-level (OFC) to low-level (PCC) processing regions - attentional control mechanism^36^. Additionally, the OFC synchronizes with anterior temporal regions in the theta band, acknowledged to work as cognitive control during WM maintenance^17^. To conclude, we demonstrated that state 1 comprises all the features matching the executive unit in Baddeley’s multicompartment model^1,37^. This state has a crucial role in attentional control, in agreement with Cowan, who identified attention as a key component of the executive unit^3^.

### State 2 – Local processing and memory storage

During and following the early stimulus encoding phase, several WM neurophysiological studies report an event-related desynchronisation (ERD) of the occipital alpha activity^11,38–40^. This effect resembles the significant deactivation of state 2 between 200-500 ms PST. This state is associated with an alpha-dominant occipitoparietal network with phase synchronization between the posterior temporal and occipital regions.

The alpha rhythm in WM dynamics is associated with modulatory activity; an increase in alpha power reflects inhibition by gating of information flow, whereas a decrease in alpha power reflects information processing^12,41,42^. The regions involved in this state were also reported to conduct letter processing and maintenance, specifically in the n-back task with visual-verbal stimuli^43^. Lochy et al. observed the left ventral occipital-temporal cortex involved in letter representations^44^. The temporal and occipital fusiform regions were also detected by Costers et al. to decrease in inhibitory alpha activity as an increase in local letter processing^12^. The suppressed inhibitory alpha in state 2 could then reflect local independent letter processing, as traditionally reported in the WM literature. However, considering the phase-coupling network of this state, the suppression of its activity could be interpreted as the desynchronization of the visual sensory network to prevent information flow to the frontal system (state 1) once the stimulus has been perceived^45^. These two interpretations are not conflicting and indicate two aspects that coexist. The description of state 2 shares several traits with the neuroimaging description of the phonological loop in Baddeley’s model, as the storage of visual-verbal stimuli^1^.

### State 3 – Sensorimotor state

With all the attentional resources allocated to encode the stimulus, other excitatory activities should be suppressed^11^. In this regard, we observe the suppression of the beta sensorimotor state, state 3, around 100 ms. The model inferred the sensorimotor state consistently with previous TDE-HMM analyses^20,22^. The spectral profile arising in the beta band is interpreted as the excitatory rhythms generated in the SMC^46^. The suppressed excitatory beta activity in state 3 prevents the ongoing encoding of WM representation from disruption^47,48^. The prolonged suppression of this beta rhythm in target than distractor trials (see Figure 6) could reflect the increased attentional and processing demand leading to the motor response in target trials^49^.

### State 5 – an M300 state

In the last phase of a WM task, the memory template is recalled, matched to the stimulus, and response selection is initiated^50,51^. These phases in the WM loop need a coordinating and integrative activity, which we assign to state 5. This state is significantly activated in the 200-500 PST time window and is characterised by a beta activity distributed in a complex spatial network including frontal, temporal, and parietal regions. The prefrontal executive regions are thought to transiently recall the memory template from the letter processing (temporal-occipital) regions and proceed with template-stimulus matching^52,53^. In alignment with this, state 5 exhibits a long-range frontotemporal phase-coupling, and its task-evoked response, as shown in Figure 5.2, peaks significantly around 350 ms in both target and distractor trials; recalling and matching memory template and stimulus occur in both conditions.

Executive frontal regions also engage in selection as part of a decision-making process ^54^. The state activation is remarkably increased and prolonged in target as compared to distractor trials (Figure 6), and this should reflect target selection and motor response preparation. This last role of state 5 is further corroborated by the involvement of the SMC and the beta activity – spectral mode 3^47^. However, as illustrated in Figure S8, a strong PSD map and phase-coupling network also form in spectral mode 2. The broadband activity of this state reflects the diversified frequency bands (theta, alpha, beta, and gamma) associated with maintenance, recall, matching, and motor plan ^32,55,56^, and the intertwined development of these processes during an n-back task. A developing research line identifies cross-frequency coupling as the underlying mechanism for WM subprocesses, explaining the broadband activity reported by traditional works ^40,56–58^. In this context, we would suggest that the broadband spectral profile of state 5 results from cross-frequency interactions.

Another typical feature associated with WM decision and matching processes is the P300 ^11,30,59^, and the temporal profile of state 5 strongly resembles the MEG equivalent, the M300 wave. While the P300 has been extensively studied in EEG literature^11,59^, its magnetic counterpart has only been recently investigated. Costers et al. identified the M300 peak in temporal regions and the SMCs ^12,14^, which are also recruited in state 5. The P300 wave has been associated with a wide broadband 25-40 Hz activity ^10,13^. Again, the same frequency range is associated with state 5.

## 5 Limitations and Future Work

As explained in the methods, the spectral content of the HMM states might be biased towards lower frequencies due to the PCA decomposition implemented prior to the model inference ^27^. Because of this, when extracting four spectral modes, we discard spectral mode 4, as it filters the noisy low-gamma activity. Nonetheless, WM dynamics have also been explored in the gamma band. While future works may investigate the gamma activity in the dynamic functional networks underpinning WM, this is missing in this work. Future studies the cross-frequency coupling within one state; this could help explain the broadband activity and the long-distance interactions, for example, in state 5.

The TDE-HMM assumes that all the inferred states are activated in a mutually exclusive fashion. However, the brain likely recruits different brain networks simultaneously. Novel analysis designs, like DYNEMO^60^, could address this limitation in the future. The TDE-HMM is a stochastic model that could provide slightly different results at every run. In our study, we ran the model several times – as explained in the results - and the inferred states showed consistent temporal, spectral, and spatial profiles across runs.

In addition to the n-back, the WM literature presents other paradigms to address working memory. Each task could yield slightly different dynamics, as they involve WM subprocesses differently ^15,48^. To validate this experimental design future works should apply the same methodology to investigate other working memory tasks (the Stenberg task or the symbol-to-digit modality test); this could help identify the task-specific from the general WM processes.

## 6 Conclusions

This study represents the first effort to unravel the multidimensional nature (time, space, and frequency) of the network dynamics underlying working memory. We used MEG data acquired during an n-back WM task and applied the TDE-HMM technique to extract data-driven dynamic functional networks^22^. The model inferred four task-relevant states with unique temporal, spatial, and spectral profiles. These provide a unified and integrated description of the characteristic traits of WM activity, which, instead, have been reported sparsely in the WM literature.

We are able to interpret the HMM states within Baddeley’s multi-component model of WM^1^. This unique exploration of WM reveals novel traits, such as the M300 state that represents a potential magnetic counterpart of the cognitive EEG P300 feature and could lead to a more in-depth understanding of cognitive processes in MEG data.

## Acknowledgment

The authors would like to thank the participants for their time and commitment to this study.

## Funding Information

The MEG data collection was enabled by grants from the Belgian Charcot Foundation and by an unrestricted research grant provided by Genzyme-Sanofi. CR is funded by Fonds Wetenschappelijk Onderzoek (FWO, Grant numbers: 11K2823N, 11K2821N).

## Conflict of Interest

The authors declare no potential conflict of interest.

## Data Availability Statement

The data for this study are not publicly available. Researchers interested in a collaboration on these data are welcome to contact the senior authors. Analysis scripts are available upon request from the corresponding author.

## Supplementary Materials

### 1. Reaction time and Accuracy of response

**Figure S1.**
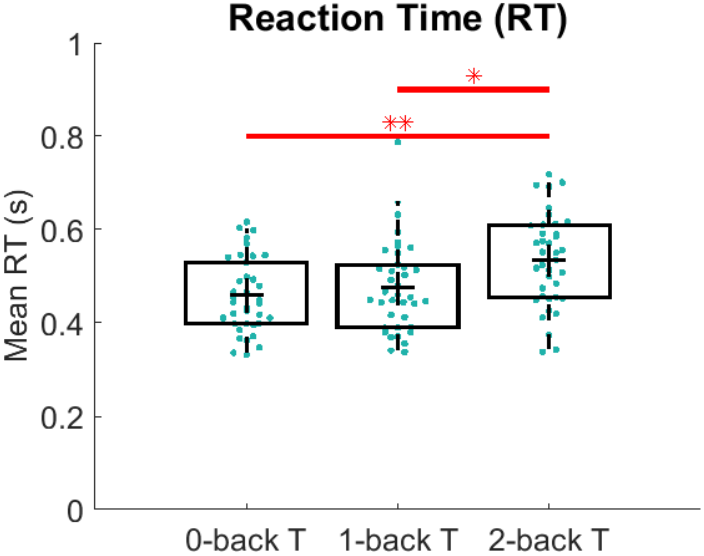
Distribution of mean reaction times (RTs) between the three paradigm target conditions (0-back, 1-back, 2-back). The mean RTs for the 2-back is significantly increased compared to the 0-back and the 1-back conditions. (Wilcoxon rank-sum test, *0.005< p <0.05, **p <0.005)

**Figure S2.**
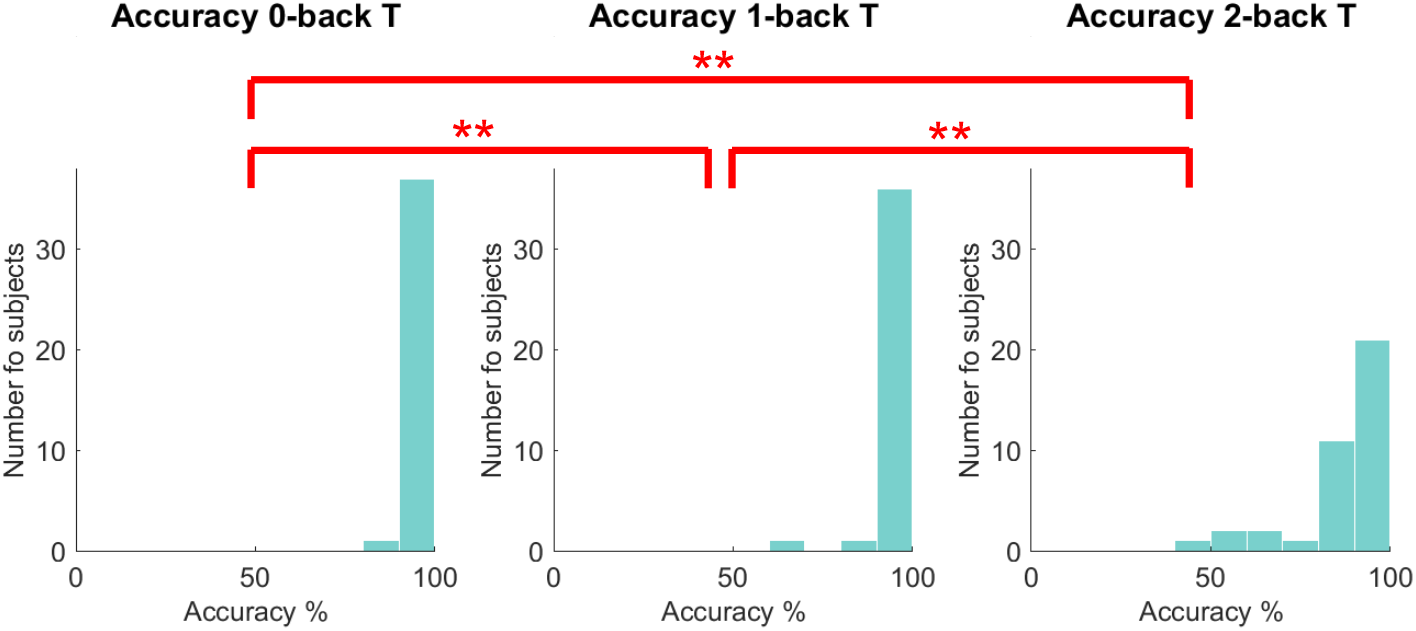
Distribution of accuracy of response for the three target conditions (0-back, 1-back, 2-back). The accuracy of response decreases significantly when increasing the WM load, from 0-back to 2-back conditions. (Wilcoxon signed rank test, *0.005< p _value<0.05, **p_value <0.005)

The mean reaction times (RTs), as the time between the stimulus onset and the button press, are 458.7±79.0 ms for 0-back T, 477.0 ±96.4 ms for 1-back T, and 534.4±98.0 ms for 2-back T. The mean RTs increase with increasing WM load (0, 1, 2-back T). In particular, the mean RTs during the 0-back T are significantly faster than the 2-back T (Wilcoxon rank-sum test, p_value<0.005) and the same hold for the 1-back T as compared to the 2-back T (Wilcoxon rank-sum test, p_value<0.05). The increasing reaction time in parallel with the increased working memory load is often reported in the WM literature and reflects the increasing task difficulty. Additionally, we explored the worsening task performance considering the response accuracy, defined as the number of correct answers over the total target trials for the specific task condition. We computed the accuracy for the 0-back T 99.3±3.3 %, for the 1-back T 97.0±6.9 %, and for the 2-back T 87.8±14.5 %. The accuracy of response decreases significantly with the increasing WM load (Wilcoxon rank-sum test, p_value<0.05).

### 2. Temporal characteristics

The temporal characteristics depict the temporal behaviour of a state over the whole model inference. These descriptors are computed starting from the Viterbi path without including any task information. The fractional occupancy (FO) measures the portion of time occupied by one state, the lifetime (LT) represents the average activation time of a state, and the interval time (IT) represents the average time window between two consecutive activations of a specific state. We refer to ^1^ for details on the computation of these quantities.

Considering the fractional occupancy (FO), states 1, 2, and 3 are activated each for about 18% of the time, and state 6 occurs 14% of the time. Instead, state 4 is activated significantly less than all the other states (Wilcoxon rank-sum test, p_value <0.005) with a FO of 11%, and state 5 occurs significantly more frequently than all the other states, with a FO of 22% (Wilcoxon rank-sum test, p_value <0.005). The average lifetime (LT) across all states is 93 ms. States 1, 2, 3, and 6 are activated for approximately 74, 84, 80, and 79 ms, respectively. Instead, states 4 and 5 show significantly longer LTs than the rest of the states: 123, and 116 ms (Wilcoxon rank-sum test, p_value<0.005), respectively. The interval time of states 1, 2, 3, and 5 are 416, 397, 383, and 506 ms, respectively. Instead, the interval time of state 4 is about 1 second, which is significantly longer than all the other states (Wilcoxon rank-sum test, p_value<0.005), and state 6 also appears less frequently than states 1, 2, 3, and 5, with an IT of 506 ms. The temporal characteristics of these states are consistent with the literature considering resting-state and task data ^1,2^. All the p values are corrected for multiple comparisons using an FDR function ^3^.

**Figure S3.**
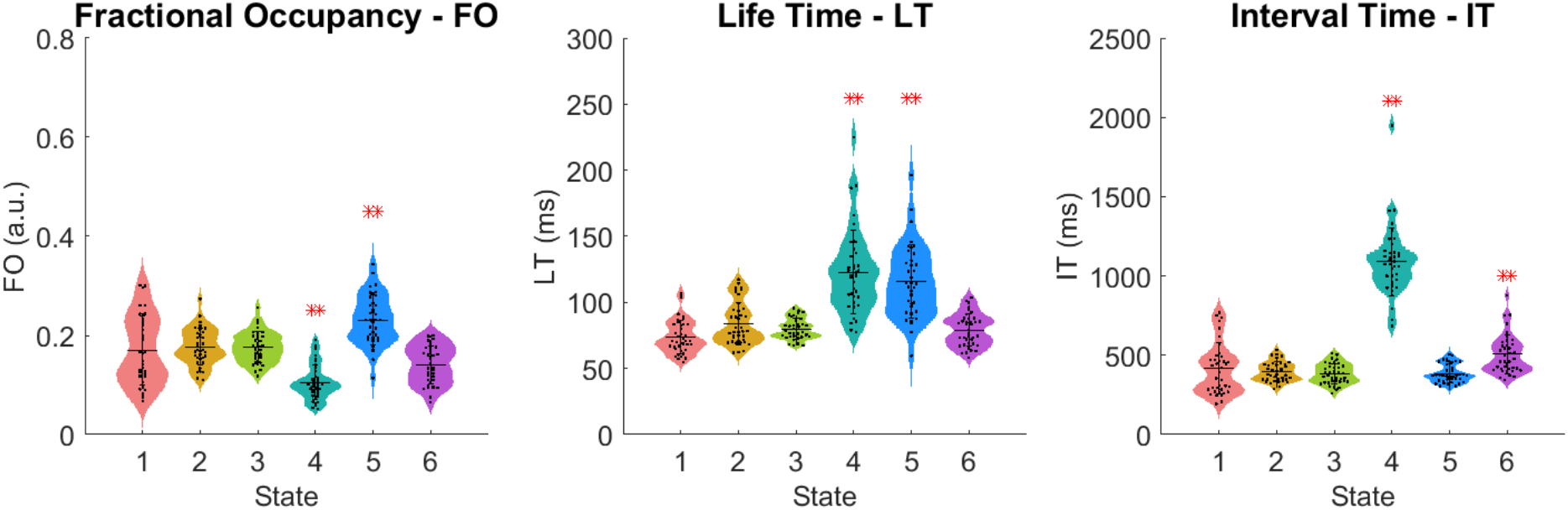
Temporal characteristics extracted from the Viterbi path of continuous data. From left, the fractional occupancy (FO) is evenly distributed across the 6 states, with an average FO per state around 16%. The life time (LT) varies across state, with very short states (1,2,3, and 6) with life times around 60 ms and long states such as 4 and 5 with a LT around 100 ms. The interval time (IT) of state 4 is over 1 sec, whereas the other states occur every 500 ms on average. ** Wilcoxon rank-sum test, p_value<0.005. All the p_values are FDR corrected to solve the multiple comparisons issue.

### 3. States Description

**Figure S4.**
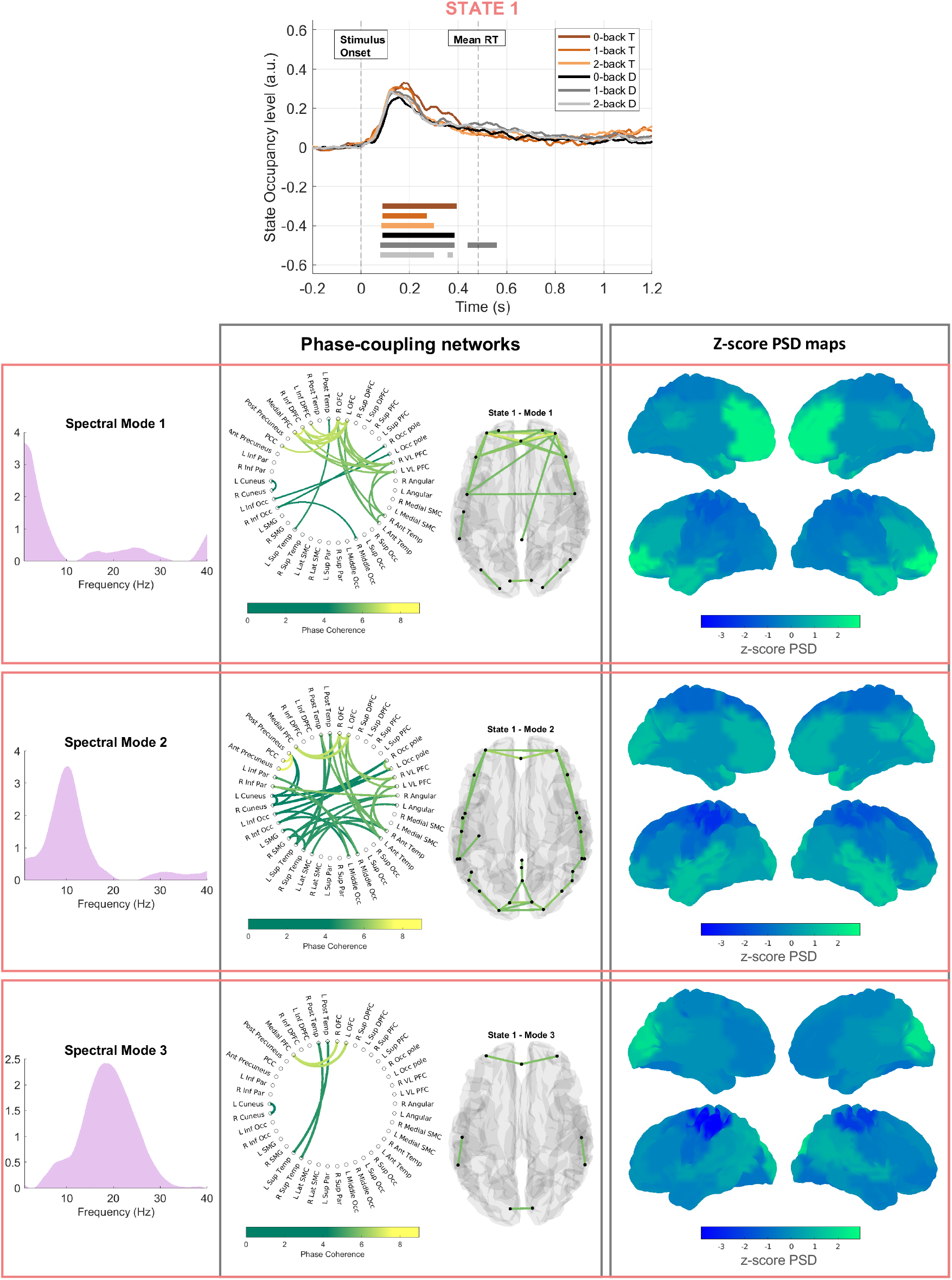
**State 1** - On top: the task-evoked occupancy level of the state for all the paradigm conditions separately. In the table, the rows consider all the profiles referred to the same spectral mode; the three spectral modes are reported in the first column. The second column shows the connectivity networks with the circular graphs and the brain glasses, and the third column shows the PSD distributions over the brain.

**Figure S5.**
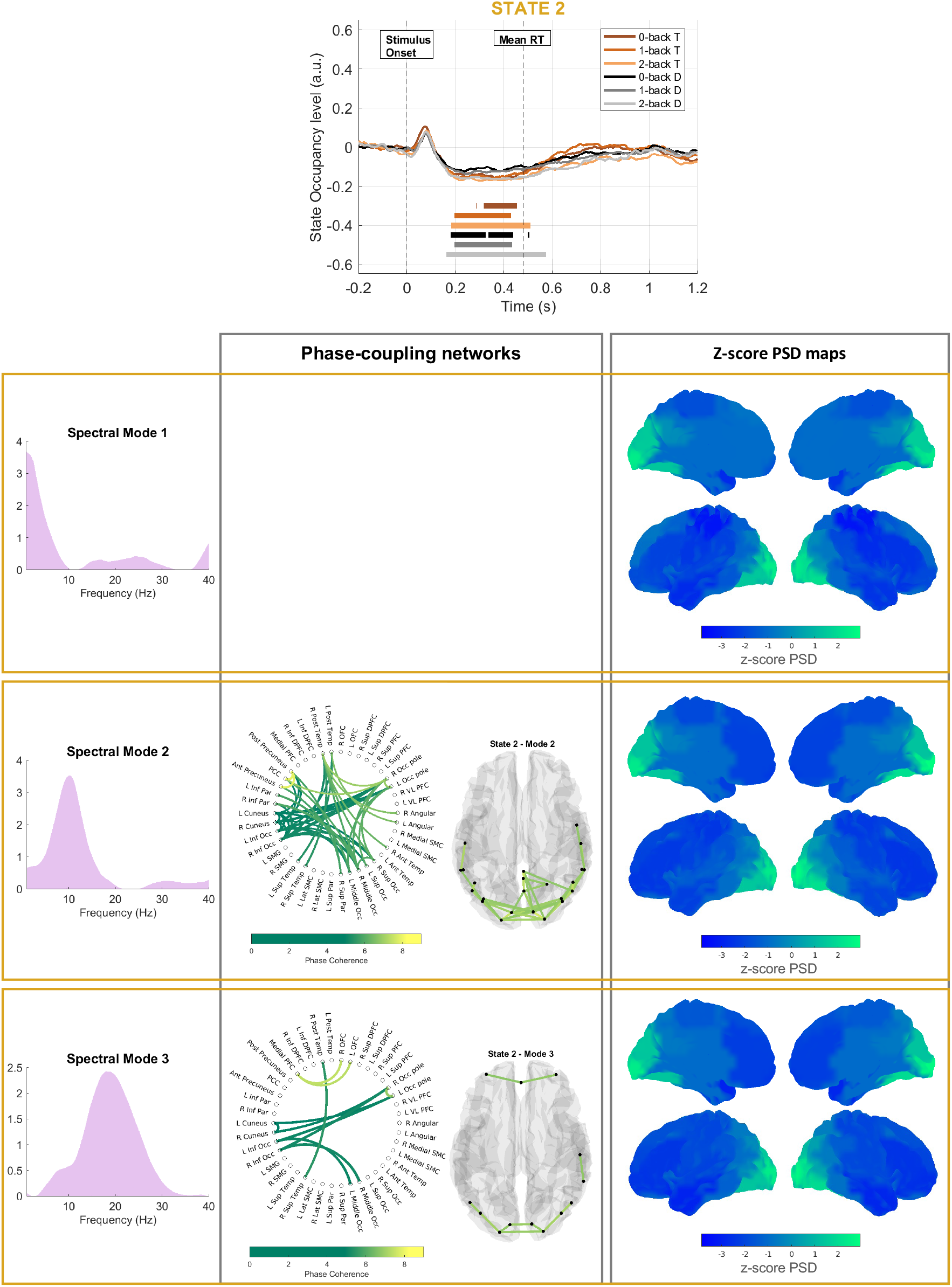
**State 2** - On top: the task-evoked occupancy level of the state for all the paradigm conditions separately. In the table, the rows consider all the profiles referred to the same spectral mode; the three spectral modes are reported in the first column. The second column shows the connectivity networks with the circular graphs and the brain glasses, and the third column shows the PSD distributions over the brain. The empty box in the connectivity networks column shows that no connections survived thresholding for the connectivity network referred to spectral mode 1.

**Figure S6.**
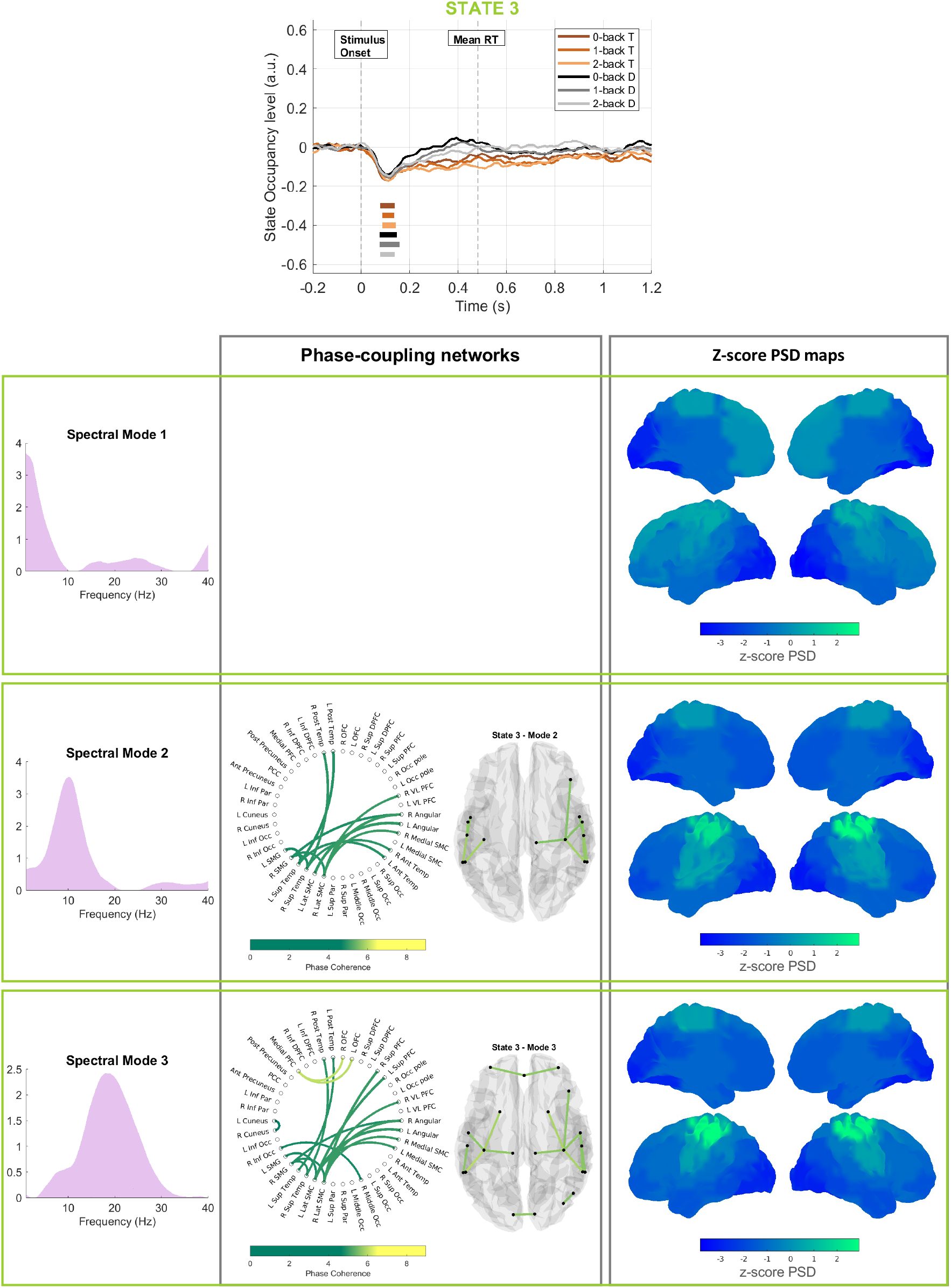
**State 3 -** On top: the task-evoked occupancy level of the state for all the paradigm conditions separately. In the table, the rows consider all the profiles referred to the same spectral mode; the three spectral modes are reported in the first column. The second column shows the connectivity networks with the circular graphs and the brain glasses, and the third column shows the PSD distributions over the brain. The empty box in the connectivity networks column shows that no connections survived thresholding for the connectivity network referred to spectral mode 1.

**Figure S7.**
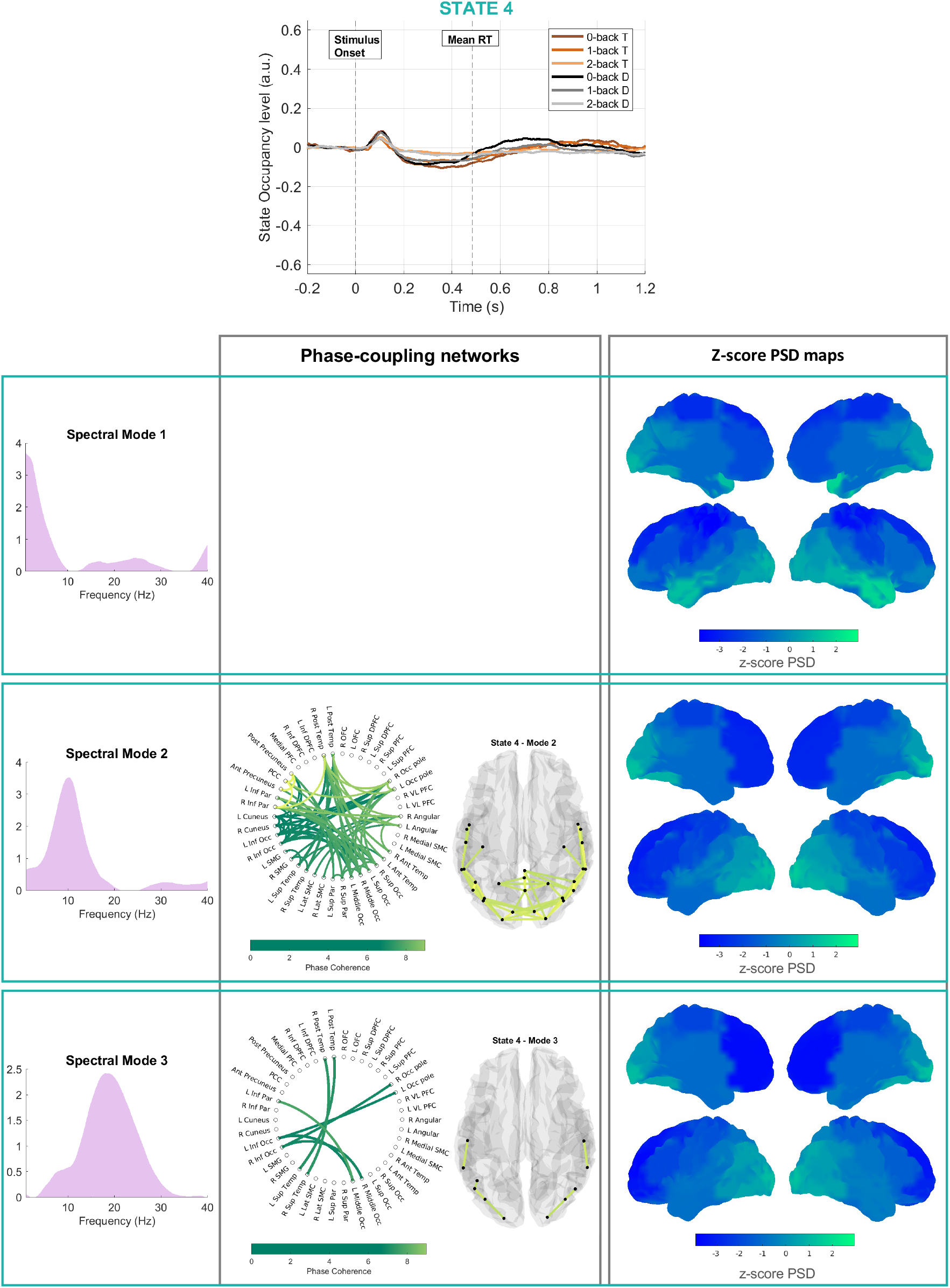
**State 4 -** On top: the task-evoked occupancy level of the state for all the paradigm conditions separately. In the table, the rows consider all the profiles referred to the same spectral mode; the three spectral modes are reported in the first column. The second column shows the connectivity networks with the circular graphs and the brain glasses, and the third column shows the PSD distributions over the brain. The empty box in the connectivity networks column shows that no connections survived thresholding for the connectivity network referred to spectral mode 1.

**Figure S8.**
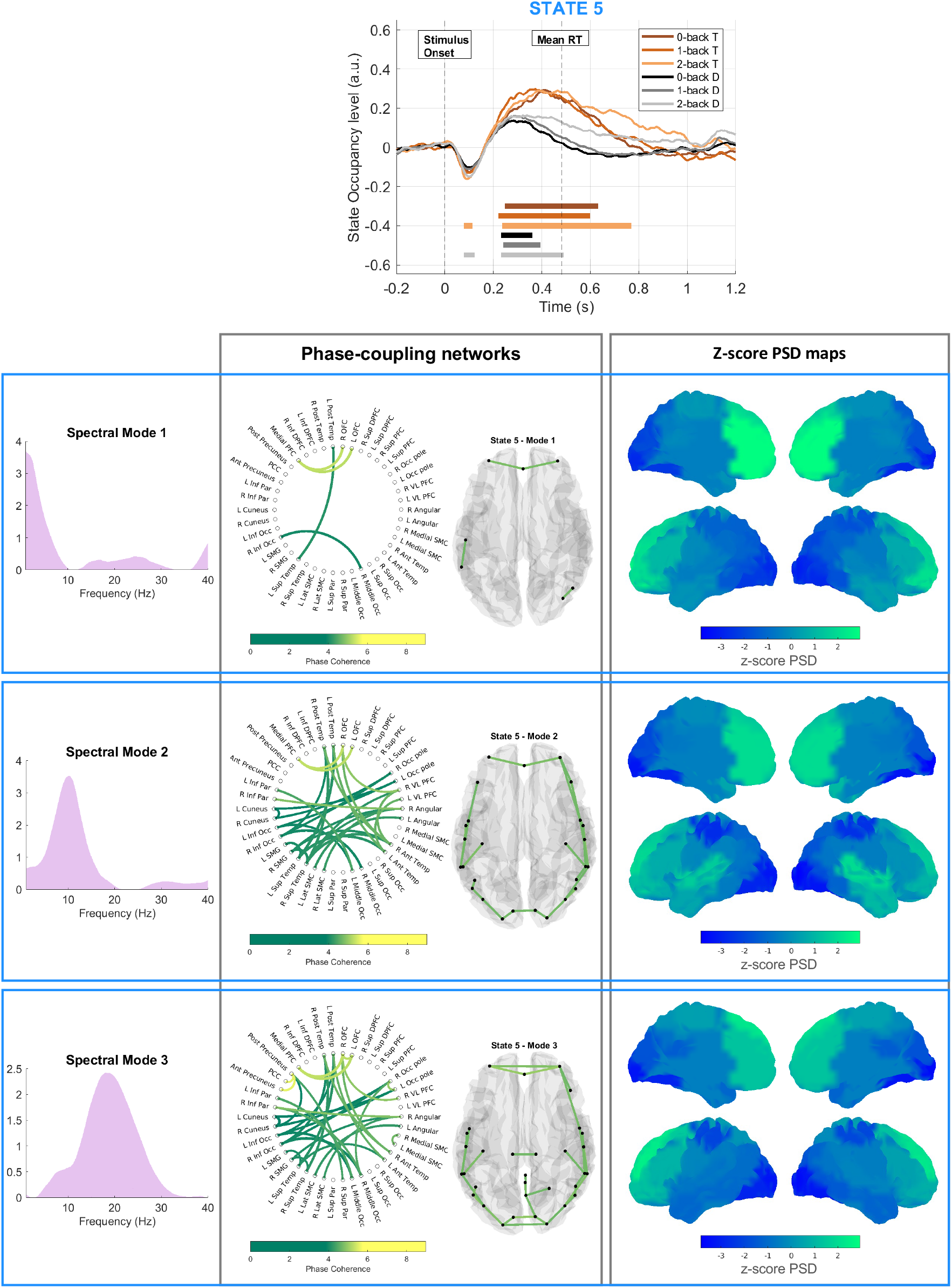
**State 5 -** On top: the task-evoked occupancy level of the state for all the paradigm conditions separately. In the table, the rows consider all the profiles referred to the same spectral mode; the three spectral modes are reported in the first column. The second column shows the connectivity networks with the circular graphs and the brain glasses, and the third column shows the PSD distributions over the brain.

**Figure S9.**
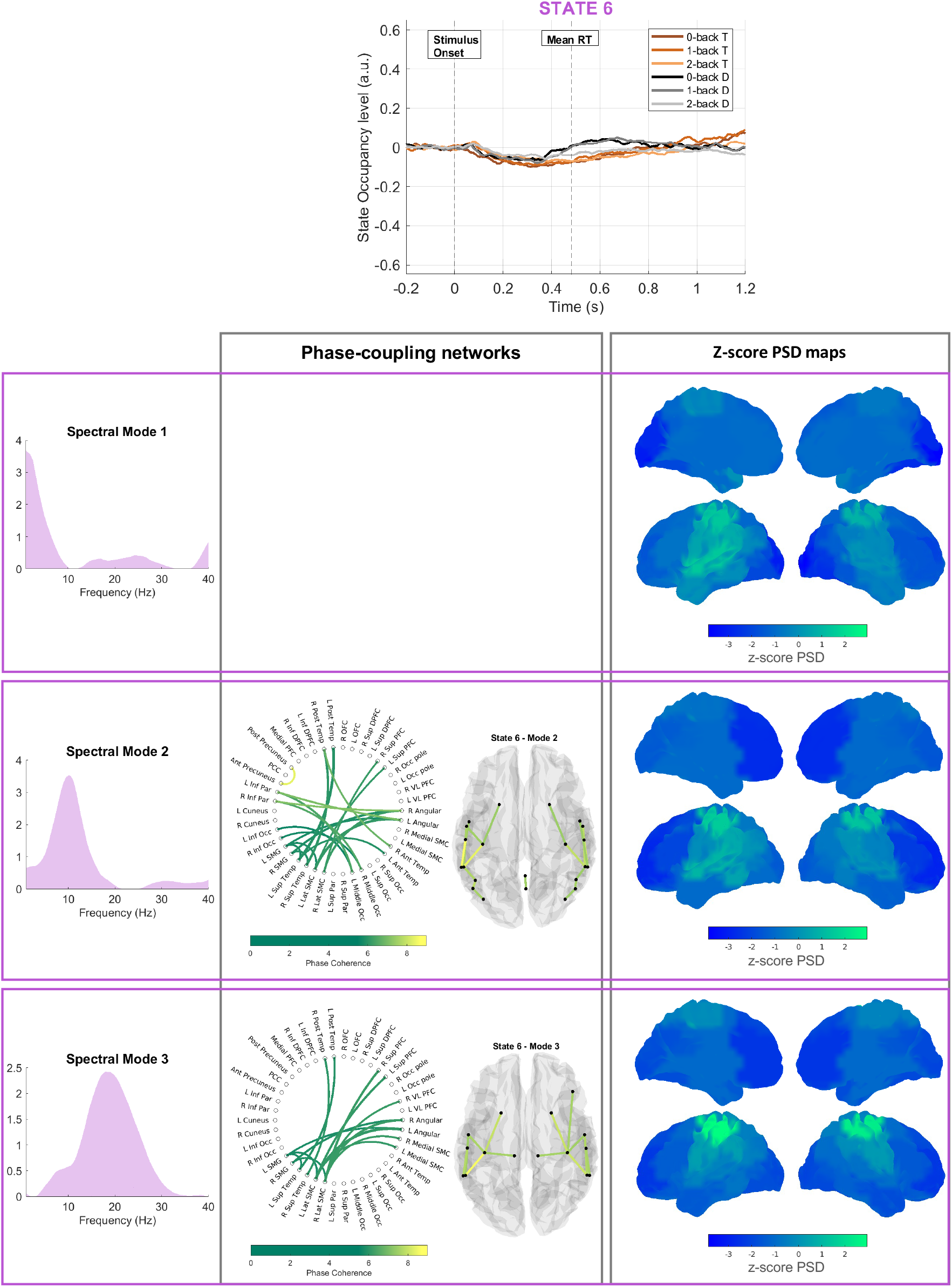
**State 6 -** On top: the task-evoked occupancy level of the state for all the paradigm conditions separately. In the table, the rows consider all the profiles referred to the same spectral mode; the three spectral modes are reported in the first column. The second column shows the connectivity networks with the circular graphs and the brain glasses, and the third column shows the PSD distributions over the brain. The empty box in the connectivity networks column shows that no connections survived thresholding for the connectivity network referred to spectral mode 1.

### 4. Targets vs Distractors: target recognition, response selection, and motor response

**Figure S10.**
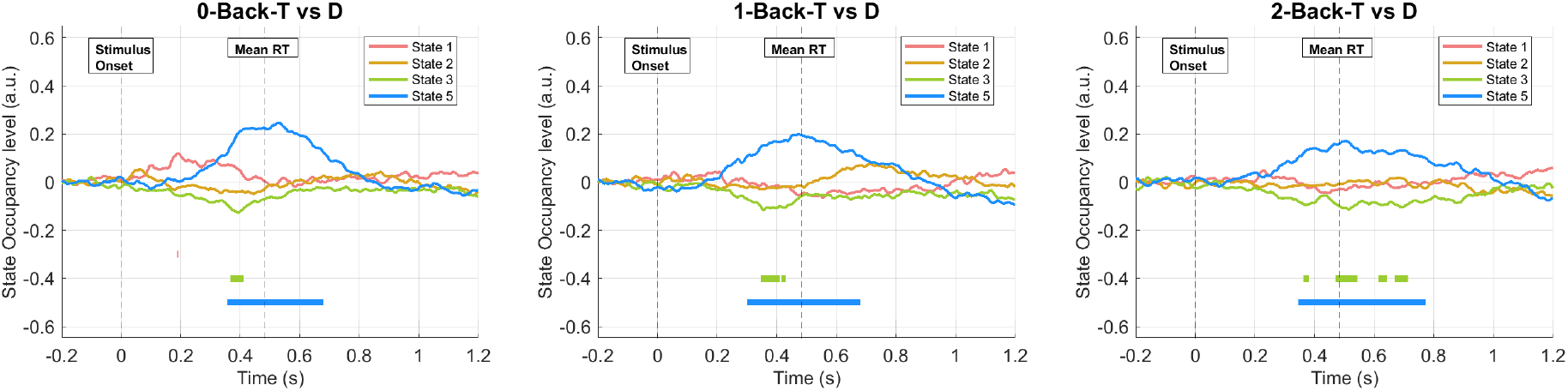
We display the contrasts between target and distractor trials for each WM load condition: 0-back, 1-back, 2-back, from left to right. State 5 shows an amplified activation in target trials as compared to distractors in all WM load conditions, which corroborates the involvement of this state in response selection/moto planning – a process that takes place in target but not in distractor trials. When visually comparing the 3 WM load conditions, we observe that the amplification effect seems to decrease from 0 to 2 back. The higher the WM load, the higher the interindividual difference in controlling strategy to perform the task ^4^. This could then lead to a smaller overall effect reported by the GLM analysis.

### 5. Scaling WM load

**Figure S11.**
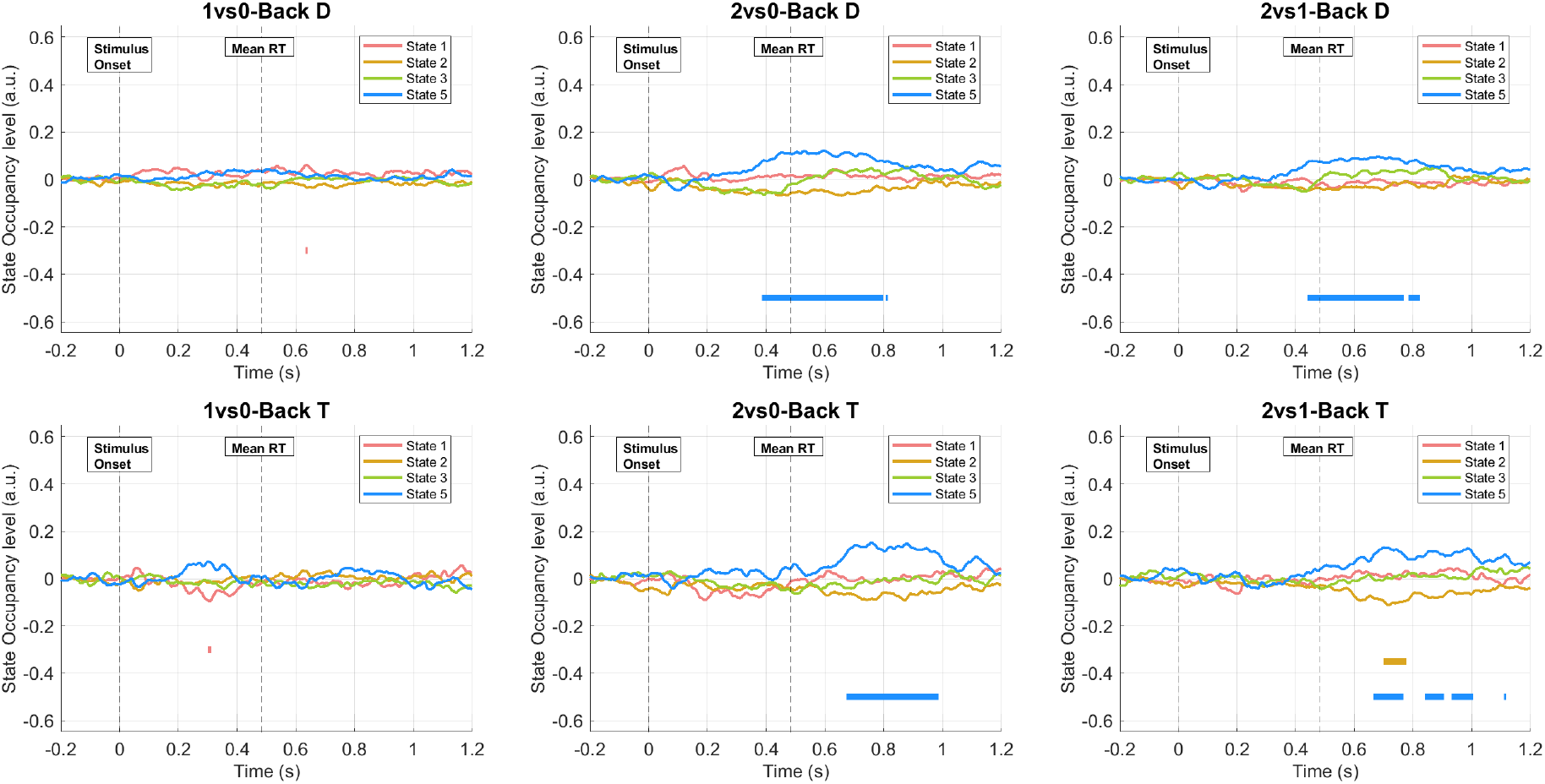
We plot the GLM contrasts of parameter estimate to investigate the effect of WM load on the task-evoked pattern of activation of the states. From left to right, we present the 1vs0-back contrast, the 2vs0-back contrast, and the 2vs1-back contrasts; the top row reports the distractor conditions, and the bottom row presents the target conditions. It is worth noticing that state 5 significantly increases in task-occupancy level in the 2 back than 0 and 1-back conditions, both in target and distractor cases.

Considering the M300 temporal wave of this state, we observe the reverse effect as compared to the electric counterpart. The P300 peak decreases with increasing WM load as an effect of attention reallocation and increasing interindividual difference in task performance strategy ^5–7^. Instead, we observe an increase M300 wave, which echoes an increase in broadband alpha-beta activity. This leads to an additional consideration; we were not able to replicate the effects of scaling WM load as reported in the MW literature: increasing theta and decreasing alpha and beta activity with increasing WM load. This point represents a limitation of this study that should be further investigated.

## Notes

### Competing Interest Statement

The authors have declared no competing interest.

