## Supplementary materials for "A novel description of the network dynamics underpinning working memory"

### 1. Reaction time and Accuracy of response

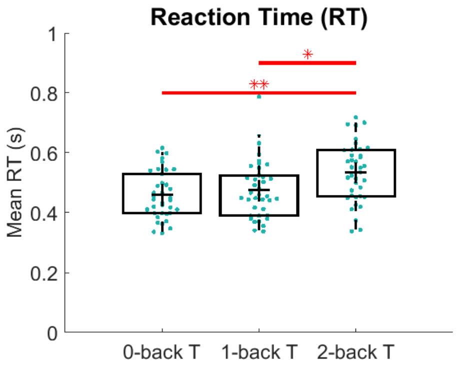


**Figure S1** Distribution of mean reaction times (RTs) between the three paradigm target conditions (0-back, 1-back, 2-back). The mean RTs for the 2-back is significantly increased compared to the 0-back and the 1-back conditions. (Wilcoxon rank-sum test, *0.005< p <0.05, **p <0.005)

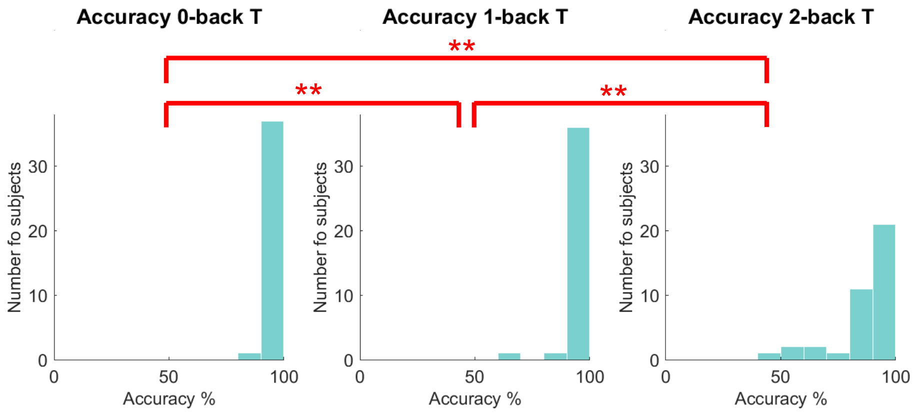


**Figure S2** Distribution of accuracy of response for the three target conditions (0-back, 1-back, 2-back). The accuracy of response decreases significantly when increasing the WM load, from 0-back to 2-back conditions. (Wilcoxon signed rank test, *0.005< p _value<0.05, **p_value <0.005)

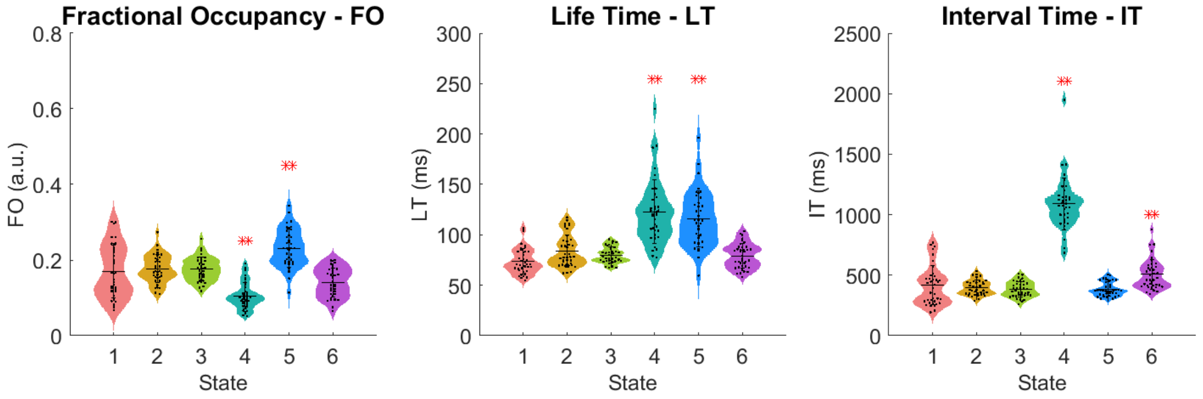


**Figure S3 Temporal characteristics extracted from the Viterbi path of continuous data.** From left, the fractional occupancy (FO) is evenly distributed across the 6 states, with an average FO per state around 16%. The life time (LT) varies across state, with very short states (1,2,3, and 6) with life times around 60 ms and long states such as 4 and 5 with a LT around 100 ms. The interval time (IT) of state 4 is over 1 sec, whereas the other states occur every 500 ms on average. ** Wilcoxon rank-sum test, p_value<0.005. All the p_values are FDR corrected to solve the multiple comparisons issue.

### 3. States Description

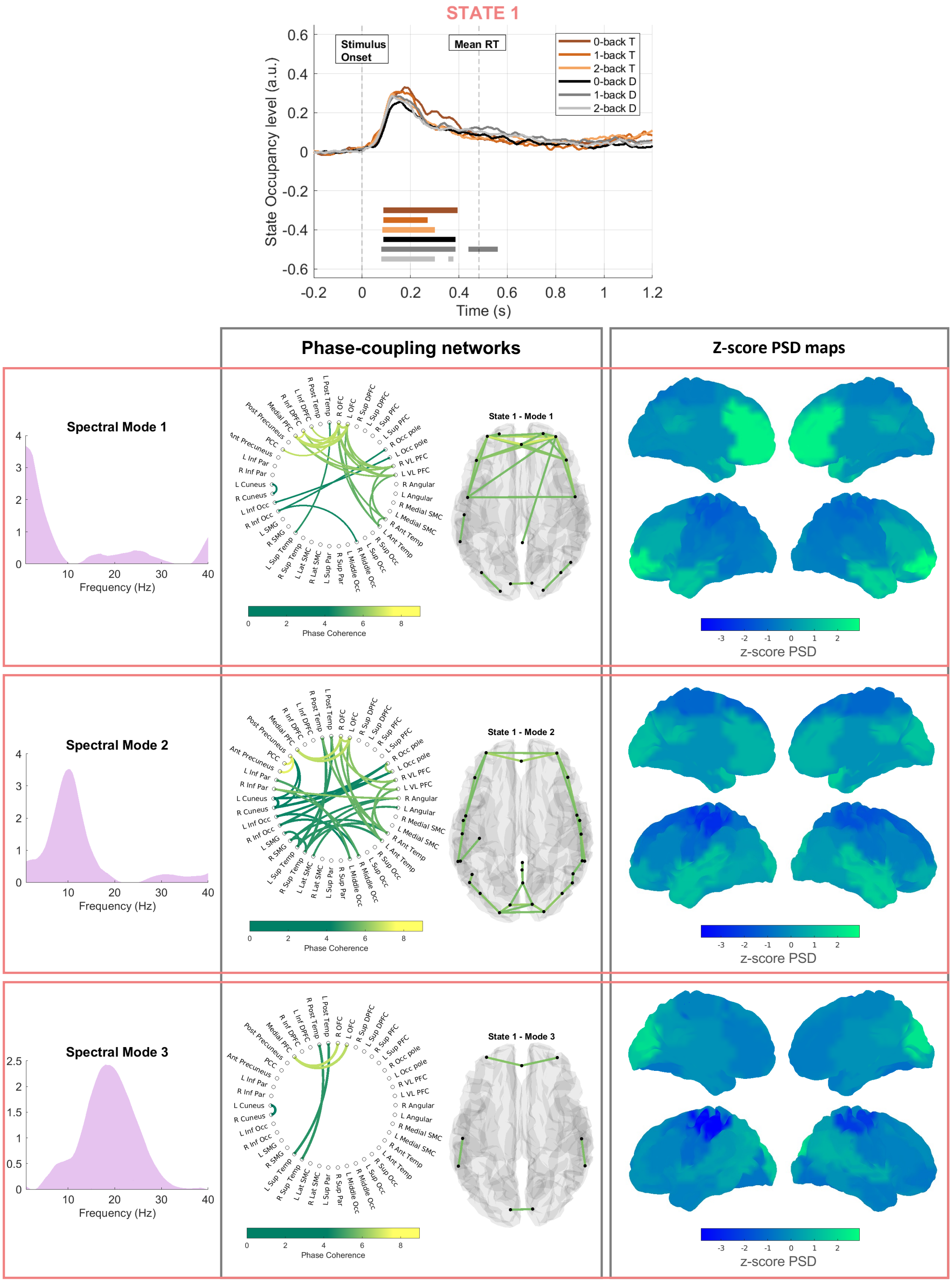


**Figure S4** – **State 1** - On top: the task-evoked occupancy level of the state for all the paradigm conditions separately. In the table, the rows consider all the profiles referred to the same spectral mode; the three spectral modes are reported in the first column. The second column shows the connectivity networks with the circular graphs and the brain glasses, and the third column shows the PSD distributions over the brain.

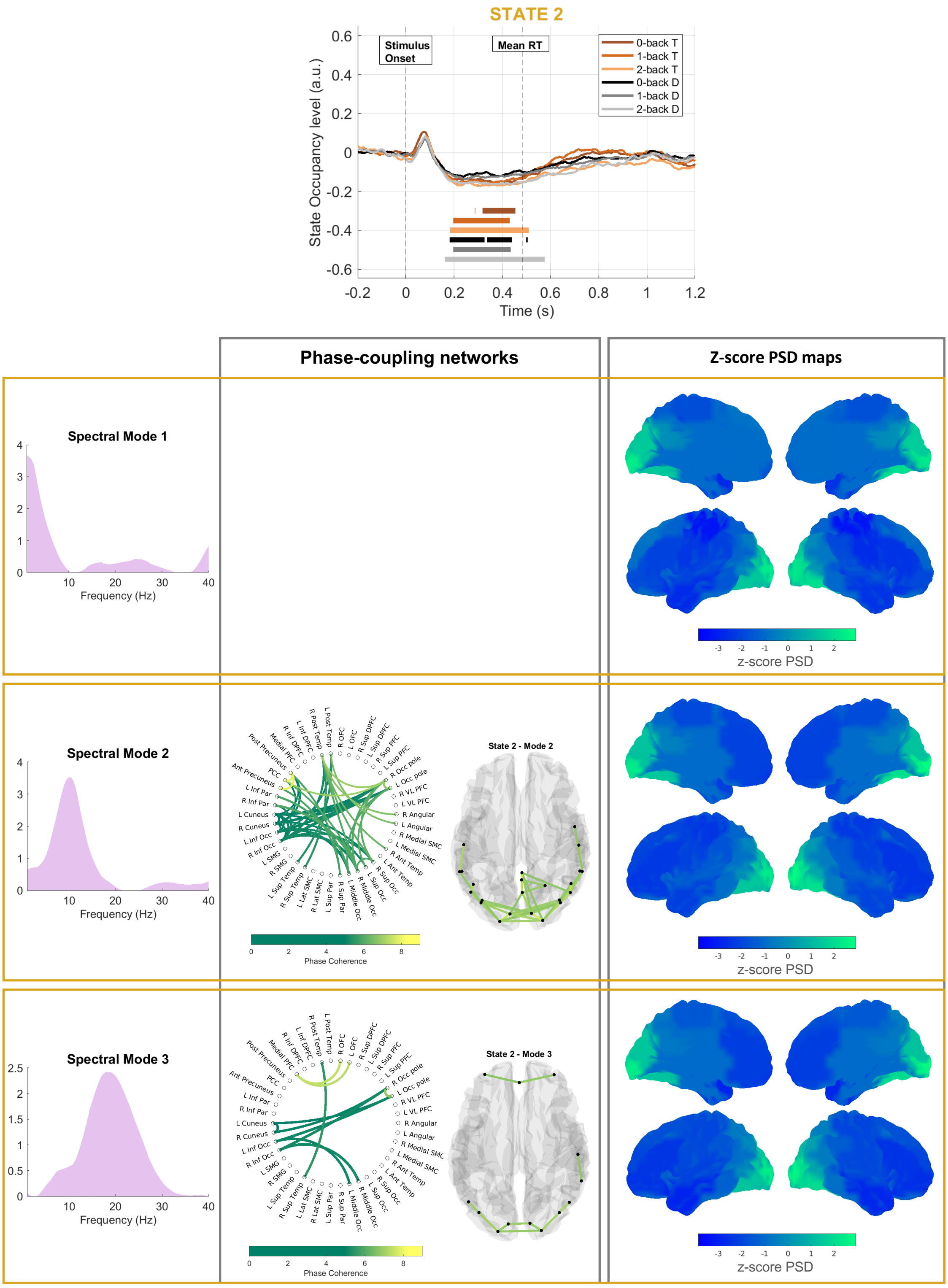


**Figure S5** – **State 2** - On top: the task-evoked occupancy level of the state for all the paradigm conditions separately. In the table, the rows consider all the profiles referred to the same spectral mode; the three spectral modes are reported in the first column. The second column shows the connectivity networks with the circular graphs and the brain glasses, and the third column shows the PSD distributions over the brain. The empty box in the connectivity networks column shows that no connections survived thresholding for the connectivity network referred to spectral mode 1.

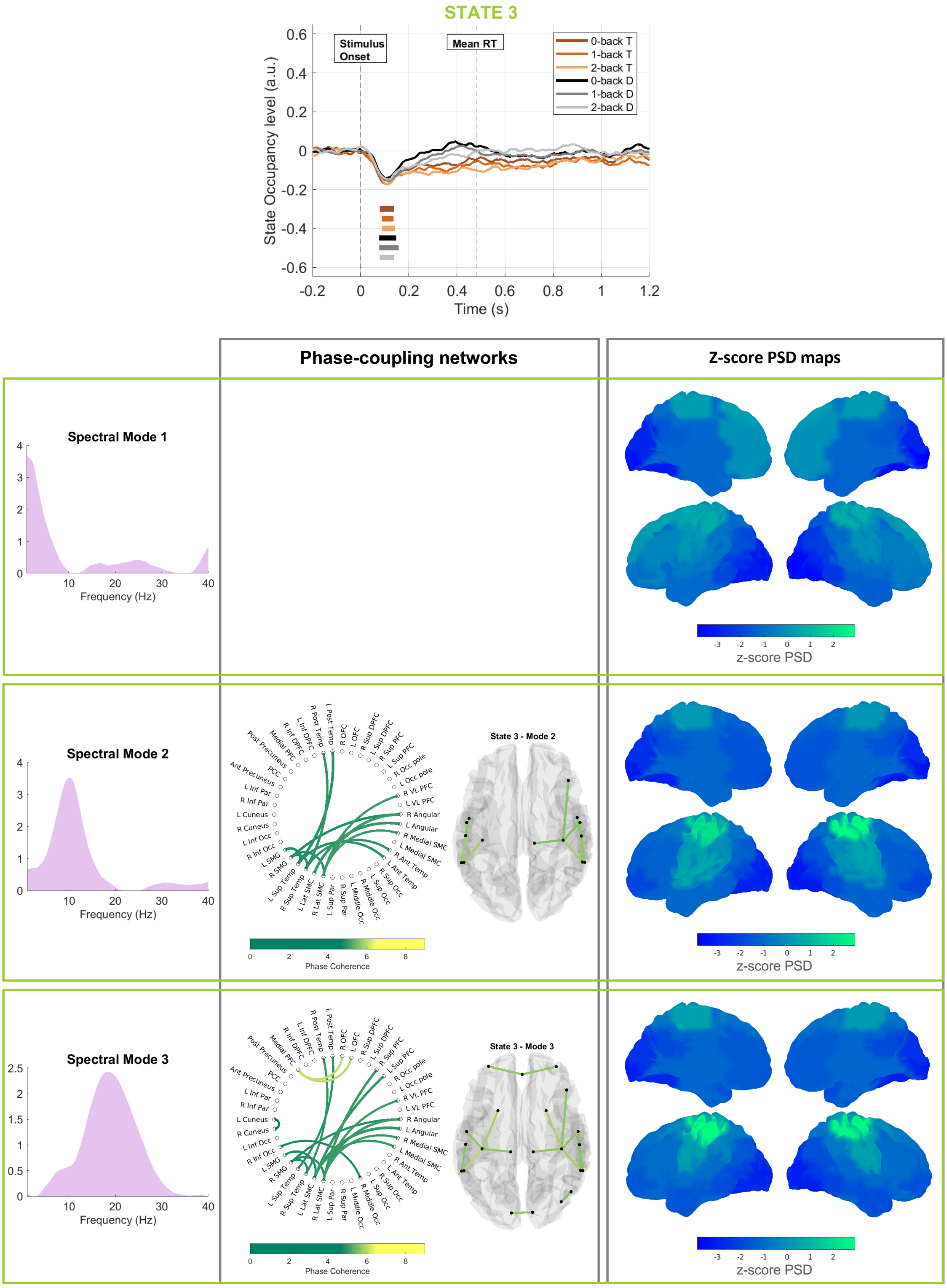


**Figure S6 – State 3 -** On top: the task-evoked occupancy level of the state for all the paradigm conditions separately. In the table, the rows consider all the profiles referred to the same spectral mode; the three spectral modes are reported in the first column. The second column shows the connectivity networks with the circular graphs and the brain glasses, and the third column shows the PSD distributions over the brain. The empty box in the connectivity networks column shows that no connections survived thresholding for the connectivity network referred to spectral mode 1.

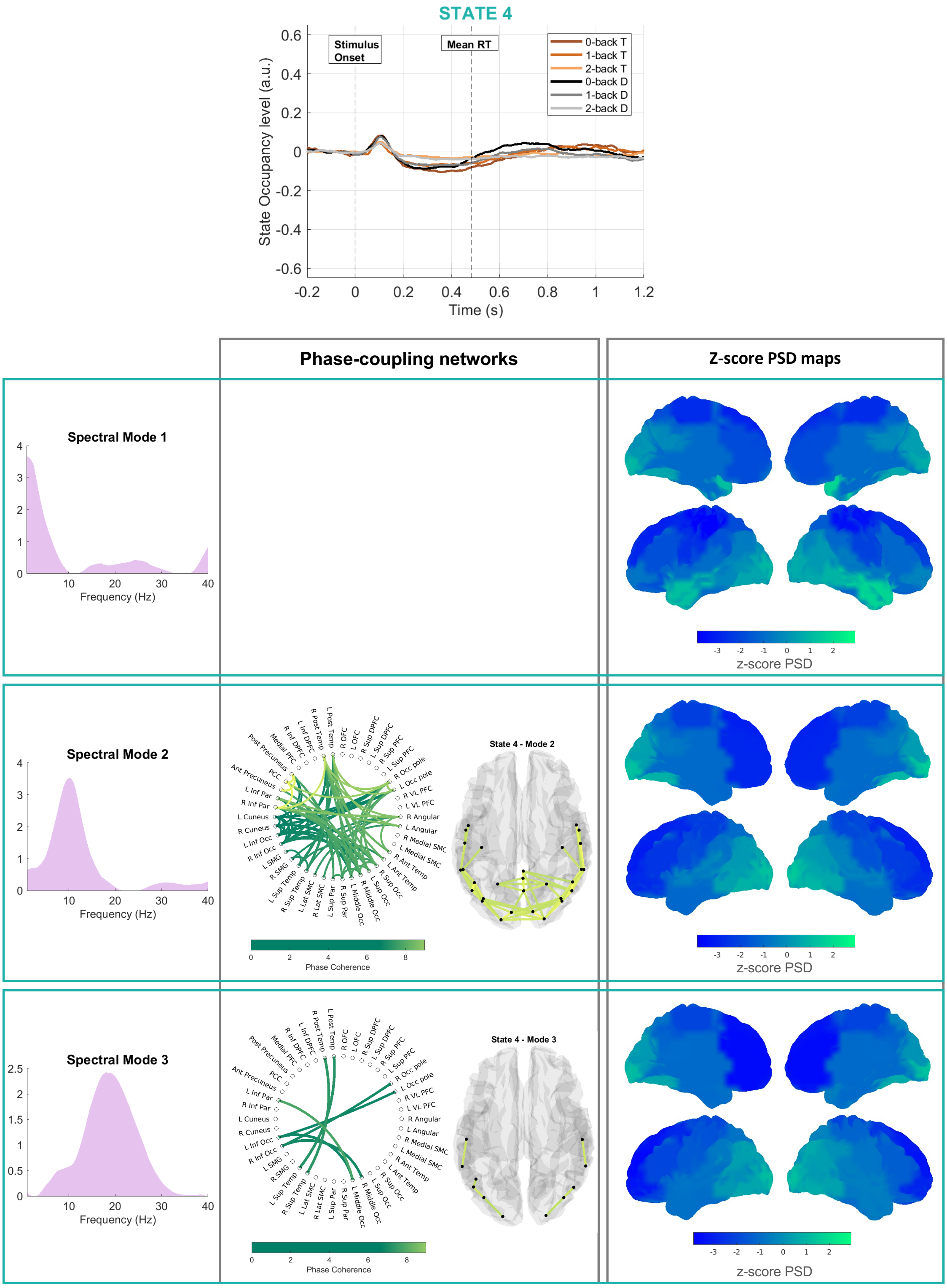


**Figure S7 – State 4 -** On top: the task-evoked occupancy level of the state for all the paradigm conditions separately. In the table, the rows consider all the profiles referred to the same spectral mode; the three spectral modes are reported in the first column. The second column shows the connectivity networks with the circular graphs and the brain glasses, and the third column shows the PSD distributions over the brain. The empty box in the connectivity networks column shows that no connections survived thresholding for the connectivity network referred to spectral mode 1.

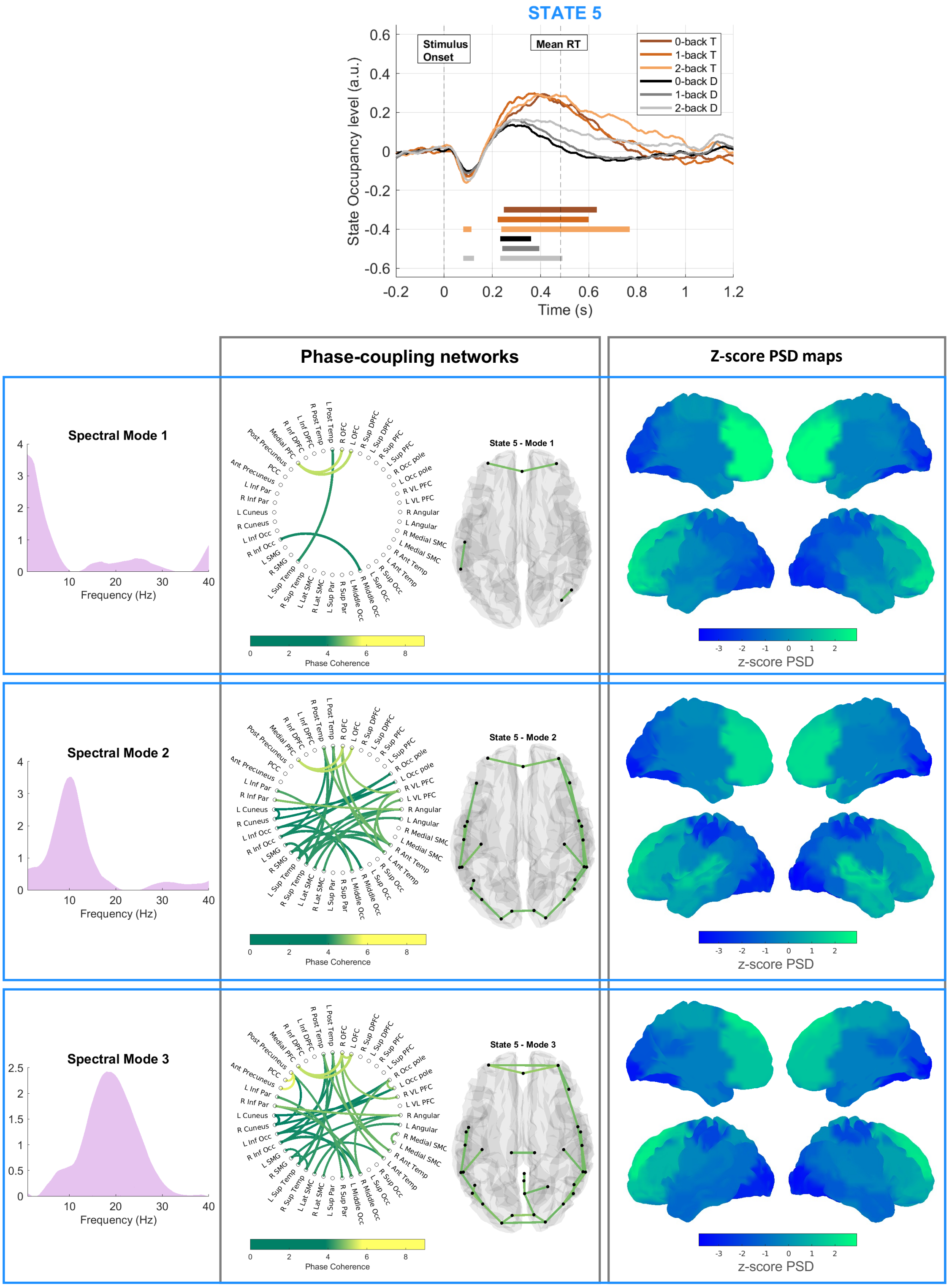


**Figure S8 – State 5 -** On top: the task-evoked occupancy level of the state for all the paradigm conditions separately. In the table, the rows consider all the profiles referred to the same spectral mode; the three spectral modes are reported in the first column. The second column shows the connectivity networks with the circular graphs and the brain glasses, and the third column shows the PSD distributions over the brain.

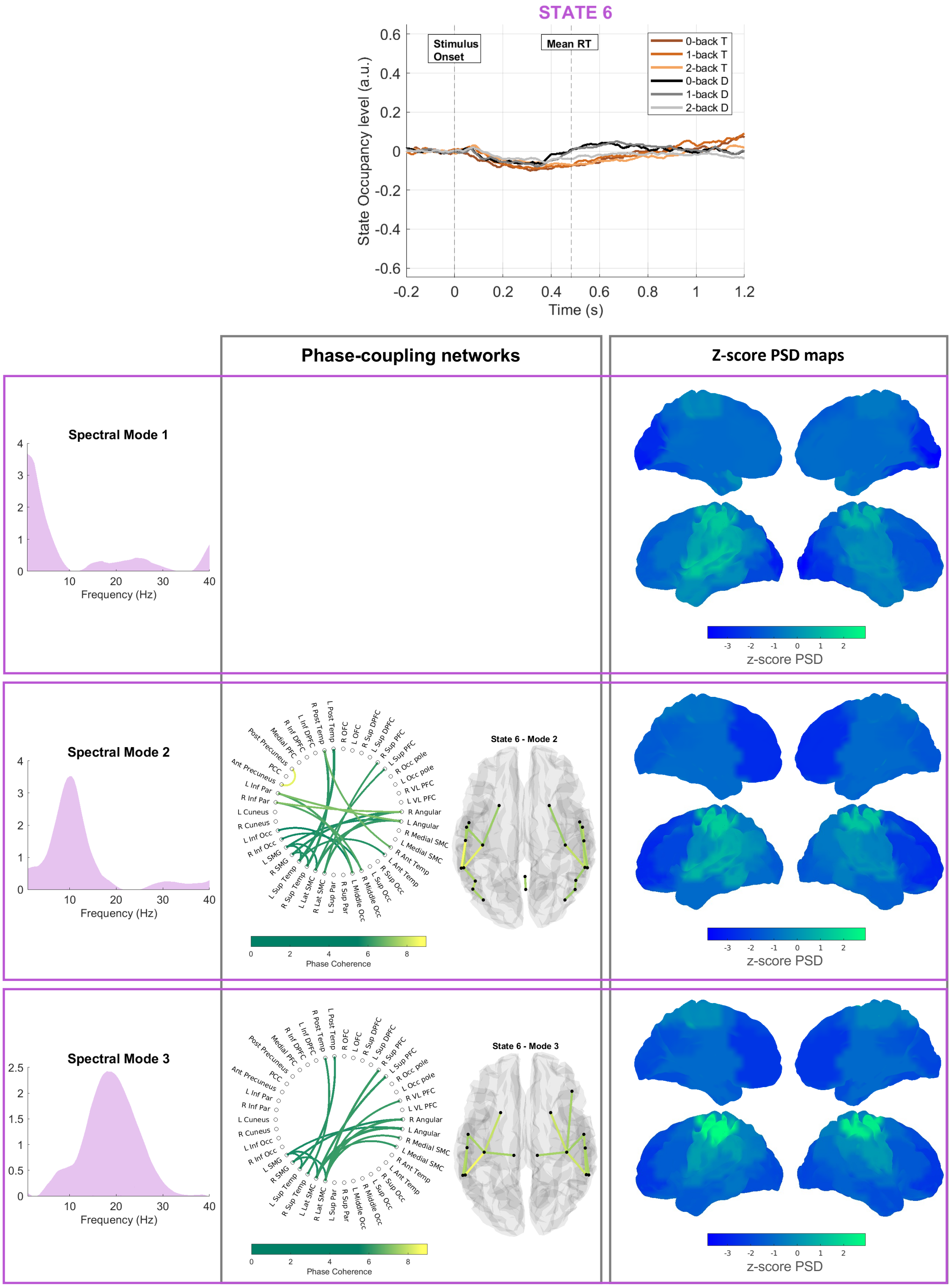


**Figure S9 – State 6 -** On top: the task-evoked occupancy level of the state for all the paradigm conditions separately. In the table, the rows consider all the profiles referred to the same spectral mode; the three spectral modes are reported in the first column. The second column shows the connectivity networks with the circular graphs and the brain glasses, and the third column shows the PSD distributions over the brain. The empty box in the connectivity networks column shows that no connections survived thresholding for the connectivity network referred to spectral mode 1.

### 4. Targets vs Distractors: target recognition, response selection, and motor response

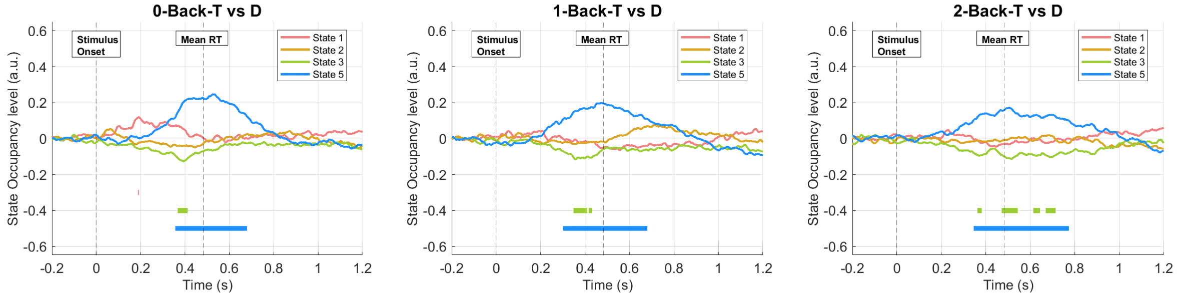


**Figure S10** We display the contrasts between target and distractor trials for each WM load condition: 0-back, 1-back, 2-back, from left to right. State 5 shows an amplified activation in target trials as compared to distractors in all WM load conditions, which corroborates the involvement of this state in response selection/moto planning – a process that takes place in target but not in distractor trials. When visually comparing the 3 WM load conditions, we observe that the amplification effect seems to decrease from 0 to 2 back. The higher the WM load, the higher the interindividual difference in controlling strategy to perform the task [4]. This could then lead to a smaller overall effect reported by the GLM analysis.

### 5. Scaling WM load

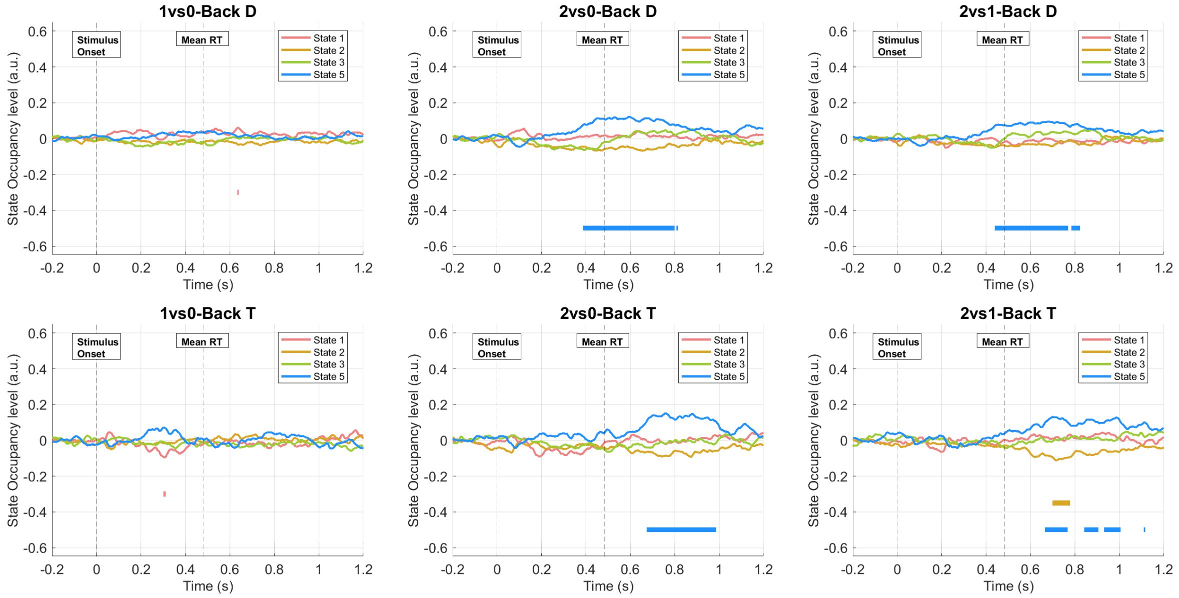


**Figure S11** We plot the GLM contrasts of parameter estimate to investigate the effect of WM load on the task-evoked pattern of activation of the states. From left to right, we present the 1vs0-back contrast, the 2vs0-back contrast, and the 2vs1-back contrasts; the top row reports the distractor conditions, and the bottom row presents the target conditions. It is worth noticing that state 5 significantly increases in task-occupancy level in the 2 back than 0 and 1-back conditions, both in target and distractor cases.

Considering the M300 temporal wave of this state, we observe the reverse effect as compared to the electric counterpart. The P300 peak decreases with increasing WM load as an effect of attention reallocation and increasing interindividual difference in task performance strategy [5–7]. Instead, we observe an increase M300 wave, which echoes an increase in broadband alpha-beta activity. This leads to an additional consideration; we were not able to replicate the effects of scaling WM load as reported in the MW literature: increasing theta and decreasing alpha and beta activity with increasing WM load. This point represents a limitation of this study that should be further investigated.
